# µFIX – Enabling Combinations of Concurrent Optogenetics and Lock-in Amplification Fiber Photometry via Removal of Optogenetic Stimulation Crosstalk

**DOI:** 10.1101/2025.02.23.639738

**Authors:** Maxim Breakstone, Spencer C. Chen, Sreya Vadapalli, Emmanuel Chavez, Lauren S. Parsonnet, Robert E. Gross, Fabio Tescarollo, David J. Barker, Hai Sun

## Abstract

Simultaneous fiber photometry and optogenetics is a powerful emerging technique for precisely studying the interactions of neuronal brain networks. However, spectral overlap between photometry and optogenetic components has severely limited the application of an all-optical approach. Due to spectral overlap, light from optogenetic stimulation saturates the photosensor and occludes photometry fluorescence, which is especially problematic in physically smaller model organism brains like mice. Here, we demonstrate the Multi-Frequency Interpolation X- talk removal algorithm (MuFIX, or µFIX) for recovering crosstalk-contaminated photometry responses recorded with lock-in amplification. µFIX exploits multi-frequency lock-in amplification by modeling the remaining uncontaminated data to interpolate across crosstalk- affected segments (R^2^ ∼ 1.0); we found that this approach accurately recovers the original photometry response after demodulation (Pearson’s *r* ∼ 1.0). When applied to crosstalk- contaminated data, µFIX recovered a photometry response closely resembling the dynamics of non-crosstalk photometry recorded simultaneously. Upon further verification using simulated and empirical data, we demonstrated that µFIX reproduces any signal that underwent simulated crosstalk contamination (*r* ∼ 1.0). We believe adopting µFIX will enable experimental designs using simultaneous fiber photometry and optogenetics that were previously not feasible due to crosstalk.

## Introduction

The human brain is responsible for reason, memory, and emotion, enabling it to control our interaction with the rest of the world. A complex network of diverse and specialized neurons performs the brain’s functions. To understand the brain is, in part, to understand the role each specialized neuron plays in this network. Modern genetic and viral engineering tools allow the targeting of genetically defined neuronal subtypes to manipulate their activity and determine their causal roles in behavior. Optogenetics is one such tool; it employs light to control ion channels (opsins) that activate or inhibit the activity of neurons^1–4^. Upon exposure to light of a specific range of wavelengths, opsins open to depolarize or hyperpolarize targeted neurons. The result of opsin manipulation or other network phenomena can be measured by imaging neuronal activity with fluorescent indicator proteins^5,6^. Some widely adopted fluorescence indicators are genetically encoded calcium indicators (GECIs; e.g., GCaMP & RCaMP variants) for monitoring intracellular calcium^7^ and genetically encoded neurotransmitter & neuromodulator indicators (GENIs)^8^. Wavelength-specific light exposure causes GECI (calcium) or GENI (neurotransmitter) binding to emit a fluorescence indicative of neural activity or inter-neuronal interaction, respectively. Whereas traditional electrophysiology indiscriminately records and stimulates all types of neurons, opsins and fluorescent indicators can be expressed in chosen cell types. This allows targeted interrogation of the neuronal network of interest, especially in understanding the mechanism underlying their pathological states.

Combining optogenetics and fiber photometry^5,6^ builds a robust methodology for real-time manipulation and simultaneous recording of targeted neurons^9–12^. However, overlaps in the spectral ranges of the available opsins and fluorescent indicators limit the experimental design of all-optical approaches. With the current GECI toolkit, it is only possible to record two fluorescence signals in the same optic fiber: one in the green (GCaMP) and one in the orange/red (RCaMP) spectra. Most readily available optogenetic tools (chiefly ChR2, C1V1, ChRmine, and ChrimsonR) overlap with one or both GECI emission spectra. In applications where multichannel fiber photometry is desired at a site nearby to the site of optogenetic stimulation, such spectral overlaps cause recording artifacts from unavoidable optogenetic stimulation crosstalk, even with high-end optical filtering or spatial separation between the optogenetic and photometry sites. Furthermore, since optogenetic stimulation usually requires at least 1 mW of stimulation power to be effective^1,2,9,13,14^, optogenetic stimulation inevitably saturates and renders nanowatt-range photometry signals unusable.

There is a lack of solutions to address crosstalk interference that still allow lock-in amplification (LIA) to capture the fiber photometry response. LIA is a common approach for recording fiber photometry^15^ by activating indicator proteins with a sinusoid-modulated excitation light intensity rather than a constant one. This produces an emitted response at the same chosen sinusoidal frequency and improves the signal-to-noise ratio. Most importantly, LIA enables encoding multiple signals in one carrier signal. Multiplexing with LIA can encode a neural signal alongside a reference isosbestic signal on the same photosensor channel, which helps correct non-neural changes in fluorescence^16–18^. However, optogenetic crosstalk interacts detrimentally with LIA demodulation, producing artifacts that mask out the photometry response during stimulation by disrupting the ability to recover the sinusoidal carrier signal.

The ability to compensate for optogenetic crosstalk would enable all-optical experimental designs that had not been previously practical due to physical proximity and spectral overlap of optogenetic and fiber photometry targets, including closed-loop experimental designs. As an example of the power of this approach, we have shown that initiating focal seizures with optogenetic stimulation in the hippocampus may involve positive feedback from the contralateral hippocampus^19^. Our work investigating re-entrant feedback in the hippocampus, under the long- standing theory that seizures arise from excitation/inhibition imbalance in neuronal activity^20^, has required readout of both the excitatory and inhibitory network activity in the contralateral hippocampus while delivering optogenetic stimulation to the ipsilateral hippocampus (see experimental setup illustrated in Fig. 1). Crosstalk has the potential to impede discoveries leveraging the combination of optogenetics and fiber photometry and removing it will open many new possibilities.

**Figure 1.**
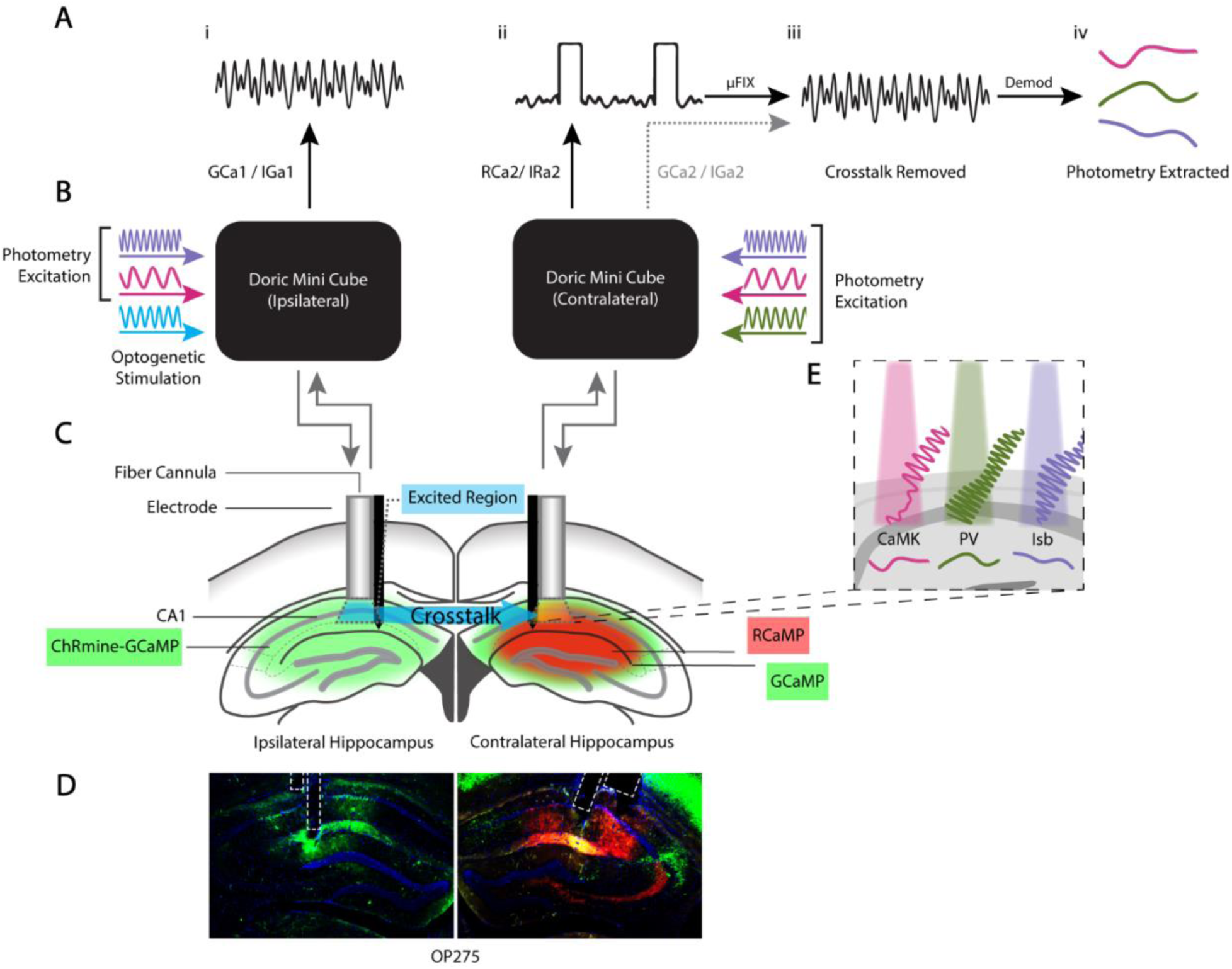
Illustration of empirical recording setup with µFIX processing pipeline. **(A)** Photosensor recordings of lock-in amplified (LIA) Ca^2+^ photometry signal without crosstalk (i) and with crosstalk (ii). µFIX removes the crosstalk from the LIA signal (iii) to recover the underlying Ca^2+^ response (iv) shown in (E). **(B)** Doric Mini Cube input/output optical filter setup converging light sources through a single fiber optic to the tissue and splitting the fluorescent emission into GCaMP and RCaMP spectra for the recording system. **(C)** Diagram of implanted components in bilateral hippocampi. Ipsilateral hippocampus: Bicistronic transduction of ChRmine and GCaMP in putative excitatory neurons. Contralateral hippocampus: PV-Cre GCaMP and putative excitatory RCaMP transduction from a viral vector cocktail. **(D)** Histology image of mouse OP275 prepared with the illustrated empirical setup. **(E)** Zoom-in visualization of the contralateral hippocampus showing the underlying LIA Ca^2+^ response.

Here, we describe the **Mu**lti-**F**requency **I**nterpolation **X**-talk (µFIX, using the Greek letter Mu) removal algorithm for recovering LIA photometry signals from crosstalk. We validated µFIX using *in vivo* recording and simulated LIA photometry signals as a ground truth. We demonstrated that µFIX introduces minimal distortion when applied to *in vivo* recordings from the mouse hippocampus and defined the extent of crosstalk that µFIX can remove. Our results show that µFIX recovers a highly accurate estimate of the underlying photometry response originally corrupted by optogenetic stimulation crosstalk. Extended simulations indicate that µFIX signal recovery is highly accurate over a robust range of stimulus durations commonly employed to activate neural activity, and neural population-level temporal dynamics of fluorescence indicators.

## Results

We prepared mice to study the excitation/inhibition imbalance during seizure generation by simultaneously monitoring Ca^2+^ fiber photometry activity in putative excitatory neurons and parvalbumin-positive (PV+) interneurons in the mouse hippocampus while inducing seizures with optogenetic stimulation. We transduced excitatory neurons in the left hippocampus with a bicistronic viral vector containing ChRmine for optogenetic stimulation and GCaMP for fiber photometry recordings in two PV-Cre mice. In the right hippocampus, we used RCaMP for excitatory neuron photometry and GCaMP for PV+ interneuron photometry. The experimental setup for these two mice is illustrated in Fig. 1. Their setup produces six photometry channels: three primary Ca^2+^ indicator signals with one corresponding isosbestic signal reference each to correct for non-neural fluorescence^21^. Optical filtering was employed using Doric mini cubes to converge the excitation light and separate the emission spectra (see Methods for optical filter makeup). We adopted LIA photometry to multiplex all of them simultaneously. Here, we describe the unavoidable optogenetic crosstalk in this experimental setup and the technique used to remove it.

### Cross-hemispherical optogenetic crosstalk between ChRmine and RCaMP

Optogenetic activation of hippocampal putative excitatory neurons induces seizures in freely moving mice ^13,14,22^. In this study, we delivered 5 ms pulse trains at 10 or 20 Hz for 30 s, observing that optogenetically-induced seizures manifest as an increase from baseline in the Ca^2+^ fiber photometry response recorded in both the ipsilateral putative excitatory (GCa1, not shown) and contralateral inhibitory neurons (GCa2, Fig. 2A, B). However, we encountered optogenetic stimulation crosstalk in the contralateral hippocampus on the RCaMP channel (RCa2) and its isosbestic reference (IRa2) (Fig. 2A, B). This occurred despite advanced optical filtering and as much as a 3.2 mm mediolateral (ML) separation between the recording and the stimulation sites (at the same anteroposterior (AP) axis and dorsoventral (DV) axis coordinates). The demodulated RCaMP signal with crosstalk contamination typically appeared as an oscillating signal pattern or a step increase or decrease from the baseline during stimulation. Artifacts in the RCaMP response coincide with each episode of stimulation (30s of stimulation every 120s, Fig. 2A), where any potential RCaMP response was obfuscated. The 589 nm wavelength laser used to activate ChRmine in the ipsilateral hippocampus overlapped the emission spectrum of RCaMP and was, therefore, picked up by its corresponding optical filter setup. We only needed ∼2 mW of light to activate ChRmine in the ipsilateral hippocampus, sufficient to saturate the RCaMP photosensor in the other hippocampus immediately.

**Figure 2.**
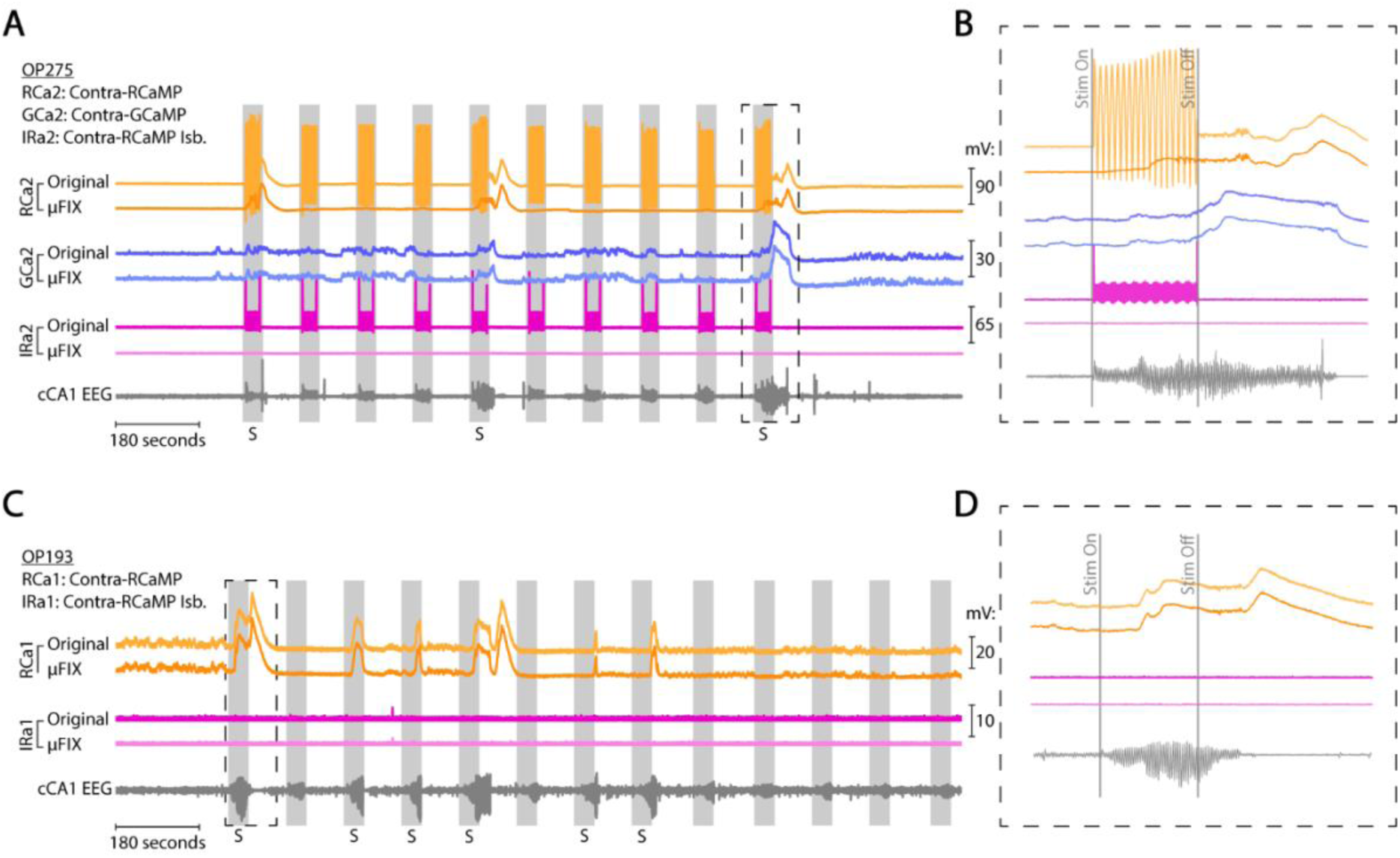
Optogenetic crosstalk and µFIX response recovery. **(A)** Photometry response before-and-after µFIX processing on a crosstalk recording. Channels RCa2 and IRa2 contain crosstalk and channel GCa2 does not. The corresponding EEG is illustrated. Gray vertical bars indicate delivery of ChRmine optogenetic stimulation, with “S” demarking a seizure response. **(B)** Zoom-in of a single seizure response epoch showing detail of recovered photometry response and EEG from (A). **(C-D)** Same as (A-B) for a recording from a mouse prepared for ChR2 stimulation. No crosstalk was observed during ChR2 optogenetic stimulation.

We explored two techniques to reduce crosstalk unsuccessfully. First, a laser light source did not remove crosstalk compared to an LED light source with the same wavelength (590nm). Although the laser light source had a narrower light beam and spectral band, neither factor was the cause for the crosstalk. Then, we postulated that light leakage occurred in the semi-translucent acrylic headcap between the dual-fiber cannula, so we adopted opaque black acrylic headcaps. This also did not reduce the crosstalk; we observed optogenetic light leaking out of the temporal side of the animal’s head during stimulation. Our investigation led us to conclude that, due to the small volume of the mouse brain, cross-cannula crosstalk came from optogenetic light reflected and diffused by the animal’s brain and skull tissue. Therefore, crosstalk was inevitable and required post-hoc processing.

### µFIX: Exploiting LIA signal composition for crosstalk restoration

LIA demodulation of the photometry signal requires a continuous sine wave without interruption or clipping; otherwise, the LIA frequency and crosstalk will interact and create artifacts. Thus, we had to address the problem within the raw photosensor recording (i.e., prior to LIA demodulation) to prevent it from disrupting signal extraction. We found that saturation of the raw recording occurred only during the delivery of an optogenetic stimulus to the tissue.

Therefore, under a 20 Hz stimulation protocol with 5 ms pulses and 45 ms pauses, there would be an intact response between every pulse for most of the time (∼45 ms). We postulated that if we can remove optogenetic artifacts by replacing these short, saturated segments with an appropriate waveform, LIA demodulation would recover the underlying photometry response. Our approach exploited the carrier frequency principle of LIA photometry – we termed it the **Mu**lti-**F**requency **I**nterpolation of **X**-talk (µFIX) algorithm.

LIA photometry works by driving the temporal profile of the fluorescence emission at the specified carrier frequencies. Our recording used 210, 330, and 530 Hz to excite RCaMP, GCaMP, and the isosbestic reference, respectively. We separated the return fluorescence into the GCaMP and RCaMP spectral ranges through a dichroic mirror and optical filter system (Doric mini cube), which are designed for precise spectral separation to ensure minimal crosstalk and to route signals to separate photosensors (Fig. 1). Power spectral analysis of the raw photosensor data from outside the stimulation period in a recording of a mouse with this experimental setup confirmed that 210, 330, and 530 Hz were the main response frequencies (Fig. 3A). However, we surprisingly observed the RCaMP 210 Hz carrier emission at the GCaMP photosensor channel and the GCaMP 330 Hz carrier emission at the RCaMP photosensor channel despite optical filtering. The 530 Hz excitation light produced both GCaMP and RCaMP isosbestic emissions; spectral peaks for all three LIA carrier frequencies were found in both RCaMP and GCaMP photosensor channels.

**Figure 3.**
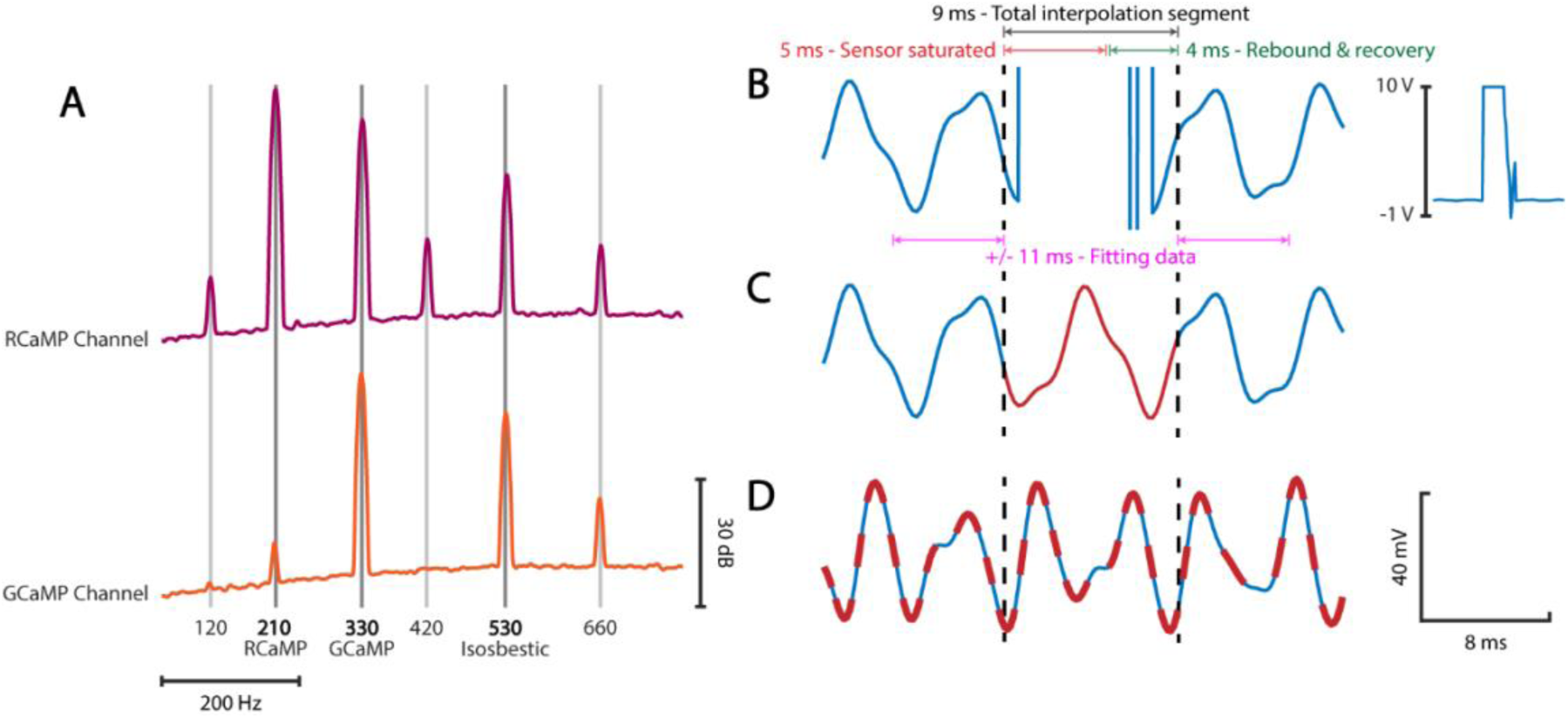
Signal component analysis and crosstalk segment fitting. **(A)** Spectral analysis of the GCaMP and RCaMP photosensor recording, showing LIA carrier frequencies used for GCaMP (330 Hz), RCaMP (210 Hz) and isosbestic reference (530 Hz). Additional peaks were found at the harmonics and beat frequencies of 210 and 330 Hz. **(B)** A segment of photosensor recording disrupted by stimulation crosstalk. Inset: same segment in (B) at the full 10 V scale. A 5ms stimulation pulse saturates the photosensor with an after-saturation artifact. A segment of 9 ms was cropped for replacement. **(C)** The same segment in (B) after µFIX interpolation (red). Interpolation was based on intact data 11 ms before and 11 ms after the cropped segment. **(D)** A different photosensor recording segment without crosstalk. µFIX was applied in the same way as if there was crosstalk. The recovered signal (dashed red line) closely matched the original (R^2^ ≈ 1.0).

We inferred that the most accurate restoration of the saturated segments would be a composite of all the LIA carrier frequencies. µFIX, therefore, works by filling in the saturated segments with interpolated patches generated using the following model of the LIA fluorescence emission at the photosensor:

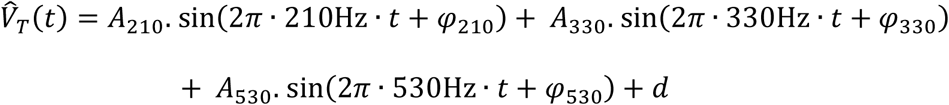

This is the expanded version of Eq(4) in the Methods for the three carrier frequencies adopted in our recording. Since GCaMP/RCaMP have slow dynamics, we make the critical assumption that the encoded fluorescence signal amplitudes (*A*_210_, *A*_330_, *A*_530_) during brief saturated crosstalk segments are constant. The amplitude (*A*_210_, *A*_330_, *A*_530_), phase (*φ*_210_, *φ*_330_, *φ*_530_), and offset (*d*) were estimated by fitting the model to the non-contaminated recording just before and after each saturated segment. For 5ms stimulation pulses, as in our experiment, we identified segments of 9 ms for µFIX – from the onset of the stimulus to 4ms after to exclude after-saturation artifacts (Fig. 3B). The model parameters were sufficiently estimated by the raw photosensor recording for 2.5 cycles of the lowest carrier frequency (11.3ms @ 210 Hz) before and after the contaminated segment (5 cycles, 22.6ms altogether).

### µFIX effectively restores the photometry response from crosstalk

We first tested whether µFIX introduces distortions to the underlying photometry response when applied to uncontaminated recordings. We found that it resulted in a perfect match to the original photosensor recording (Fig. 3D). We calculated the R^2^ value of the µFIX filled-in segments for the uncontaminated GCa2 channel plotted in Fig. 2A, finding that it was between 0.98 and 1.00 across the 5990 pulse segments in this recording, with a median of 1.00. Our result implied that LIA demodulation of the complete µFIX response matches the originally demodulated photometry response. We measured the fidelity of the signal recovery by calculating the correlation coefficient of the µFIX output to the original across each stimulation epoch, starting 2 seconds before to 4 seconds after the 30-second stimulus pulse train. The fidelity of the 10 epochs in Fig. 2A ranged from 0.98 to 1.00, with a median of 1.00.

We then applied µFIX to our crosstalk-contaminated recordings. µFIX produces interpolated segments that are visually continuous with the unaffected part of the photosensor recording between each segment (Fig. 3B). After LIA demodulation, µFIX-treated output was consistent with the photometry responses from non-contaminated channels (Fig. 2A & B, RCa2 vs GCa2): a slowly evolving Ca^2+^ signal in the recovered photometry response corresponding to the optogenetically-induced seizure. Crosstalk was successfully removed to reveal the underlying neural seizure and non-seizure responses in all 23 recordings, totaling 210 epochs of stimulation from two mice (OP2718 and OP275; Table 1). More examples of crosstalk-contaminated recordings and the recovered signals are shown in Fig. S1. These recovered responses are part of the empirical data pool used in simulations described later.

**Table 1.**
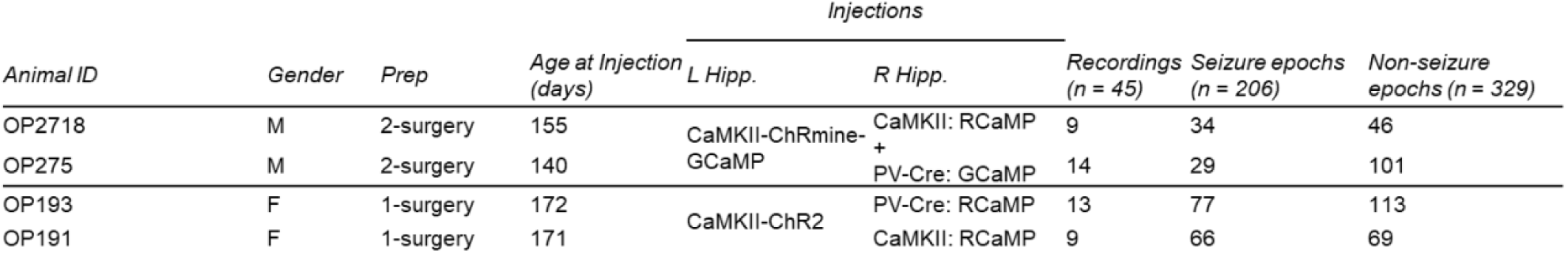
Mouse experiment parameters. Data was collected from four mice in total. Note that OP27x mice have identical setups, while the OP19x mice are identical except for right hippocampal injections.

We further verified the dynamics of the recovered photometry response by preparing two additional mice with viral combinations that did not lead to optogenetic crosstalk. These mice were transduced with ChR2 (instead of ChRmine) for excitatory neuron optogenetics in the left hippocampus, and RCaMP on its own for Ca^2+^ photometry of the right hippocampus, transduced in excitatory neurons in one mouse and PV+ interneurons in the other (Table 1). The photometry response of a recording from the latter was plotted in Fig. 2C and D. We found that the µFIX recovered RCa2 response in Fig. 2A & B exhibited similar dynamics as the uncontaminated response from the same group of neurons captured via RCa1 in Fig. 2C & D.

### µFIX recovery of artificially generated crosstalk data with high fidelity

Given the recovery fidelity of uncontaminated recordings, we inferred that the missing LIA fluorescence signal was correctly recovered for each crosstalk-saturated segment. However, since the crosstalk overwrote the original fluorescence emission signals of contaminated recordings, quantifying the effectiveness of µFIX recovery with true crosstalk was impossible. Therefore, we validated µFIX against artificially generated data via a simulated LIA photometry pipeline.

In our tests, ground truth data we assigned to our testing pipeline (the first step in Fig. 4A) served in place of unknown physiological signaling values that we sought to recover from recordings with crosstalk (starting from ii. in Fig. 1A). Unlike physiological signaling values, which were not known prior to LIA-encoded photosensor pickup and therefore lost to crosstalk, the initial value of ground truth assigned to testing was saved prior to LIA encoding and crosstalk contamination. Therefore, ground truth in our testing algorithm could be quantifiably compared before and after decoding to determine recovery fidelity.

**Figure 4.**
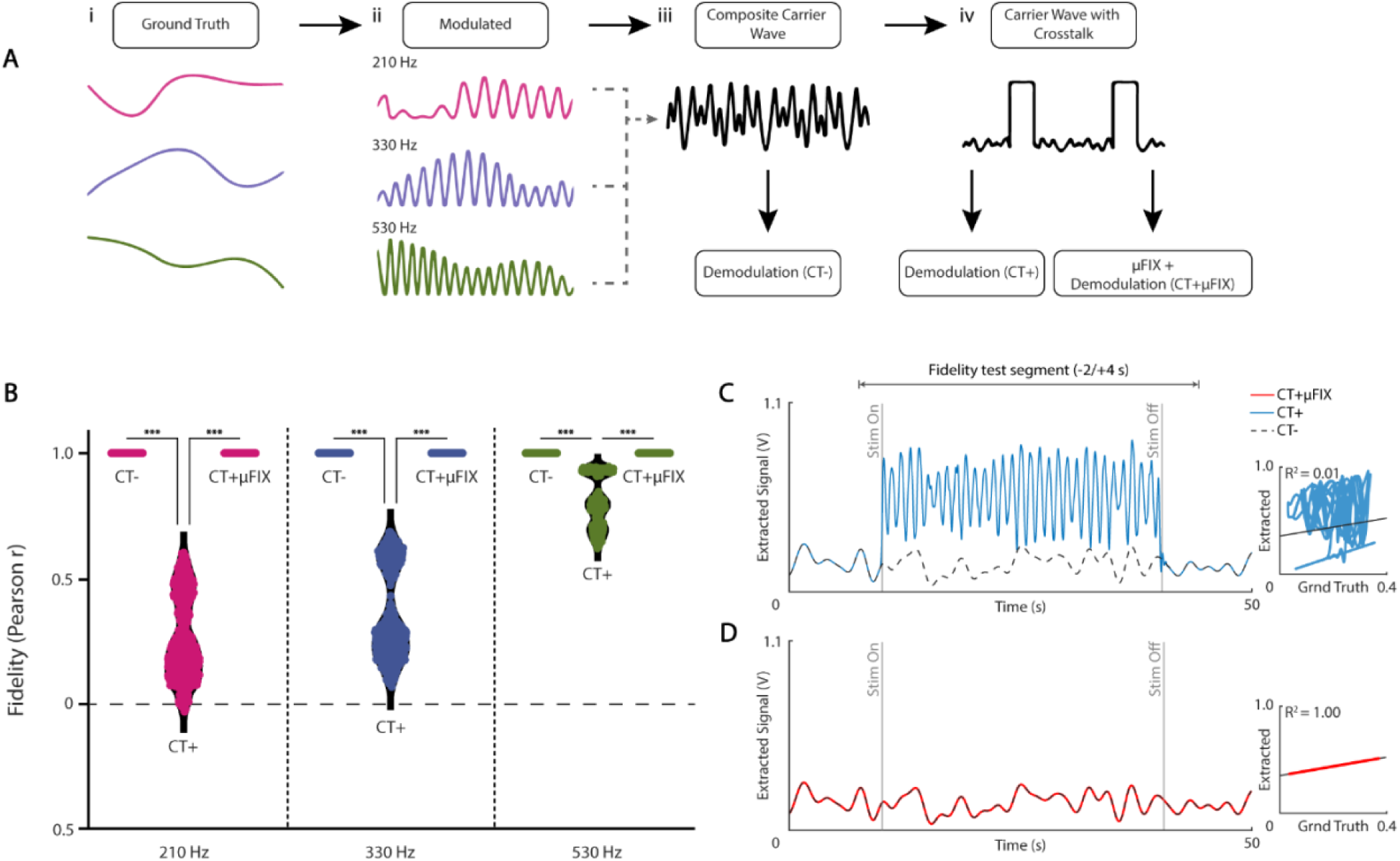
µFIX validation using simulated LIA with artificial ground truth. **(A)** Flowchart illustrating LIA photometry simulation pipeline. The ground truth signals (i) were LIA modulated (ii) and multiplexed into a single composite carrier wave (iii). Crosstalk was added to the composite carrier wave (iv). **(B)** Pearson correlation was used to compare the original ground truth to the LIA demodulated responses before simulated crosstalk (CT−), after simulated crosstalk (CT+), and simulated crosstalk with µFIX applied (CT+µFIX). Results show 120 runs of the artificial ground truth simulation in (A). In each run, three ground truth signals were generated, and the simulation was run three times with signals rotated through all three carrier frequencies. Altogether, there are n = 360 matched data points for each group. **(C**) An example epoch of demodulated response CT+ (blue) compared against CT− (dashed black line). The fidelity of demodulation was measured using Pearson’s correlation coefficient over a period from 2 s before the start of stimulation to 4 s after the end of stimulation (gray lines). Right: Scatter plot showing lack of correlation between CT+ from (C, blue) and its corresponding ground truth. By contrast, CT− strongly correlates to the ground truth (black line). **(D)** The demodulated response CT+µFIX for the same epoch as (C). Right: Scatter plot demonstrating r ≈ 1.00 correlation between CT+µFIX in (D, red) and its corresponding ground truth.

We contaminated ground truth data with artificial crosstalk and tested the effectiveness of µFIX recovery of the ground truth (Fig. 4A). We performed this validation first with artificially generated ground truth. Then, we verified it further using empirical data as the ground truth derived from a combination of uncontaminated and contaminated, µFIX-recovered recordings.

LIA photometry was generated by modulating the excitation light intensity in a sinusoidal profile (*EX*_*f*_(*t*)) with a baseline offset. The excitation light probes the underlying Ca^2+^ indicator signal (*V*_*s*_(*t*)) multiplicatively, producing a fluorescence emission (*V*_*e*_(*t*)), modelled as (see also Eq(1) in methods):

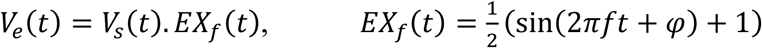

Here, we modeled the excitation light (*EX*_*f*_ (*t*)) as a sinusoid oscillating between zero and one at the carrier frequency *f*. For simplicity, we omitted the baseline offset in the excitation light.

When multiple LIA light sources are combined (multiplexed) into a single optic fiber, it can be modeled as a summation of the individual emissions (also see Eq(2) of Methods):

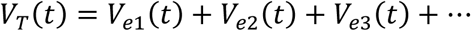

Our artificial known ground truth signals were generated with distinct amplitudes and dynamics, like empirical recordings (see Methods). We randomly generated three source signals for each simulation run, LIA-modulated them at the same carrier frequencies as our experiments (210, 330, and 530 Hz), and summed them to simulate the raw photosensor recording with all three signals multiplexed together (Fig. 4A). Rotating each triplet of generated source signals through different assignments to the three carrier frequencies allowed us to isolate source-specific effects from frequency effects; this created three data points per trial. For each trial, we first demodulated the summed emission without added crosstalk, referring to this as the CT– result.

This created a baseline measure of recovery by calculating the Pearson *r* correlation coefficient between the demodulation output and the known ground truth—a measure referred to as just fidelity from here forward. As expected, the CT– output perfectly reconstructed the ground truth with a median fidelity of 1.00 and a 95^th^ percentile range of 1.00 to 1.00 (Fig. 4B).

Therefore, we added simulated crosstalk by overwriting the ground truth carrier wave with a pulse pattern matching real crosstalk. Each stimulation pulse time was set to the saturation point of the photosensor at 10V, followed by a rebound to -1V (Fig. 3B). We added stimulation pulse crosstalk at the regular 20 Hz interval in our real recordings. This closely replicates the pattern that real crosstalk interference created in the carrier wave.

Introducing simulated crosstalk created illegible interruptions to the resultant demodulated signal (CT+), akin to the effect of crosstalk from empirical experiments (Fig. 4C). Crosstalk corrupted LIA demodulation, resulting in a low fidelity of 0.45 [0.06–0.94] (Fig 4B), significantly lower than CT– (p < 0.001, ANOVA, Tukey’s post-hoc test). Notably, the 530 Hz frequency fidelity (0.80 [0.64–0.94]) was significantly more resistant to crosstalk than 210 Hz (0.21 [0.02–0.56], p < 0.001 against 530Hz) and 330 Hz (0.28 [0.11–0.66], p < 0.001 against 530Hz), with 330 Hz being the least resistant (p < 0.001 against 210Hz, ANOVA, Tukey’s post-hoc test).

Applying µFIX (CT+µFIX) accurately restored the crosstalk-contaminated signal (Fig. 4D), achieving perfect 1.00 [1.00–1.00] fidelity for all three component frequencies (Fig 4B). This was significantly better than demodulation without removal (p < 0.001 against CT+, ANOVA, Tukey’s post-hoc test).

The distinct amplitude limits for our randomly generated ground truth data were chosen deliberately to test how multiplexed data of different sources might be affected by crosstalk differently. The fidelity of signal recovery appears to depend on the range of the signals rather than the magnitude of the signal. Ground truths with a signal range of 15–20 mV produced similar CT+µFIX fidelity (6.3 [5.5–6.5]) in Fisher Z units as signals of 5–10 mV (6.2 [5.8–7.2]), where the signal magnitude was reduced while the range was maintained. By contrast, ground truths with the same minimum magnitude but a larger signal range of 15–30 mV produced higher CT+µFIX fidelity (7.2 [7.3–7.5]). These minor differences translated to *r* = 1.00 fidelity in Pearson correlation units.

We further altered the dynamics of our artificial ground truth values to examine the limits of LIA and µFIX recovery (Fig. 5A). Signal dynamics of the encoded fiber photometry data determine how slowly or quickly the value changes up or down. For bulk neural activity recordings, such as fiber photometry, signals are usually slow changing (< 8 Hz). Signal dynamics is implemented in our artificial ground truth signals in nodes per second (nps), i.e., the number of randomly generated values per second from which the higher-sampling rate signal was smoothly interpolated. For the preceding µFIX fidelity validation with artificial ground truth, signals were generated at 3 nps based on our intended signal of interest from empirical recordings (Fig. S2A and C). The larger research community may be interested in faster signal content, so we repeated our simulations for artificial ground truth signals generated at higher nps. We found that LIA modulation (CT–) of 210 and 330 Hz began to fall below 0.99 recovery fidelity above 100 nps, and 530 Hz modulation maintained 0.99 fidelity to > 100 nps (Fig. 5B). CT+µFIX fidelity fell below 0.99 sooner, at 50 nps for 210 and 330 Hz modulation and 70 nps for 530 Hz modulation (Fig. 5C). By examining the power spectrum of the generated signals at these nps, we estimate µFIX can recover signal dynamics up to 20 Hz from crosstalk contamination (Fig. S2F). This is much higher than the signal content of our empirical recording (Fig. S2) and for bulk fluorescence indicator recordings at large.

**Figure 5.**
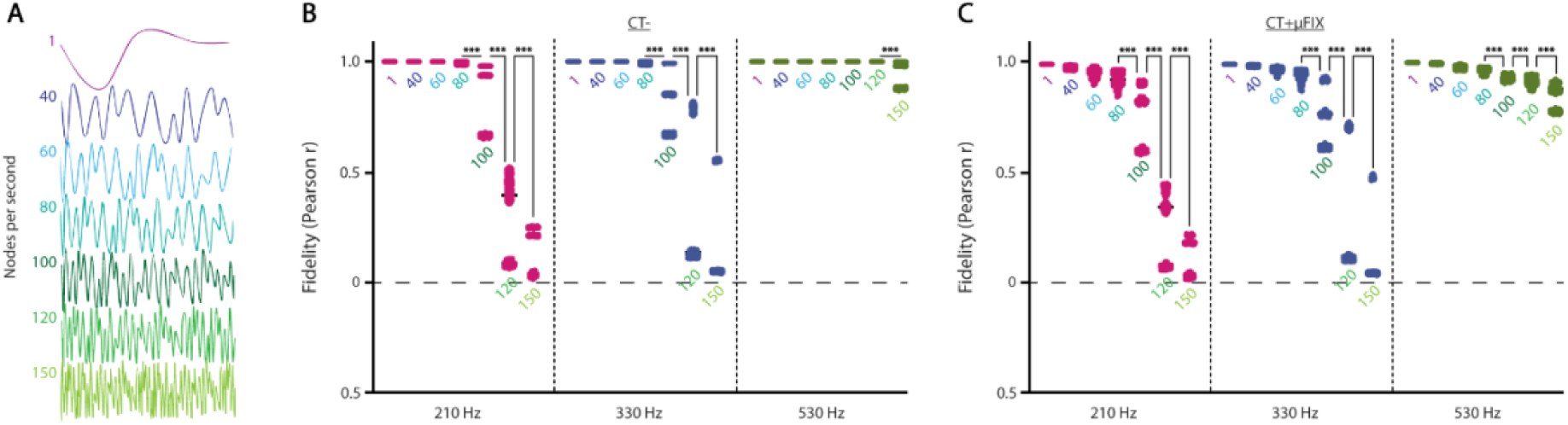
µFIX crosstalk recovery with variable source signal dynamics. **(A)** Illustration of increasing artificial ground truth signal dynamics with higher nodes per second (nps) values. **(B)** Pearson correlations by carrier frequency between the demodulated, uncontaminated response CT- (as described in Figure 4A) and the ground truth. Besides change in nps values, simulations ran as described in Figure 4. Altogether, there are n = 600 matched data points for each group. **(C)** Same approach as (B) with crosstalk added and removed (CT+µFIX).

### µFIX recovery of empirically based ground truth with high fidelity

To verify that µFIX minimally distorts real photometry recordings, we assigned them as ground truth in the crosstalk simulation pipeline. We pooled 206 seizure epochs and 329 non-seizure epochs across 45 optogenetic seizure induction recordings from four mice. Simulations were performed using the same approach for artificial ground truth and were substituted with empirical photometry epochs. We intentionally included in this pool some epochs from crosstalk-contaminated experiments to compare the result of removal of real and artificial crosstalk. All source recordings were treated with µFIX prior to use in testing for consistency. Therefore, no original crosstalk remained in the ground truth assigned to the pipeline. As with artificial ground truth, we multiplexed three empirical ground truth signals into a signal LIA simulated photosensor recording, then performed demodulation to re-extract them.

Our test indicated that without crosstalk (CT–), LIA photometry accurately captures the empirical response for all three frequencies (1.00 [0.88–1.00]) (Fig. 6C). With the addition of simulated crosstalk (CT+), the signal is unrecognizable after LIA demodulation (Fig. 6A) with low fidelity to the ground truth (0.06 [-0.28–0.88]). CT+ results were significantly lower than CT– for all frequencies (p < 0.001, ANOVA, Tukey’s post-hoc test). 530 Hz was significantly more resistant to simulated crosstalk than the other frequencies in our artificial ground truth testing (p < 0.001, ANOVA, Tukey’s post-hoc test vs. 210 and 330 Hz). After crosstalk removal with µFIX, the demodulated response (CT+µFIX) closely resembles the ground truth (1.00 [0.88–1.00]) (Fig. 6B). This significantly improved over CT+ (p < 0.001, ANOVA, Tukey’s post-hoc test). Frequency contributed a much smaller effect on the outcome for empirical recordings than in our simulated tests, based on ANOVA (Table 2, *F* = 3.1633, *p* = 0.043).

**Figure 6.**
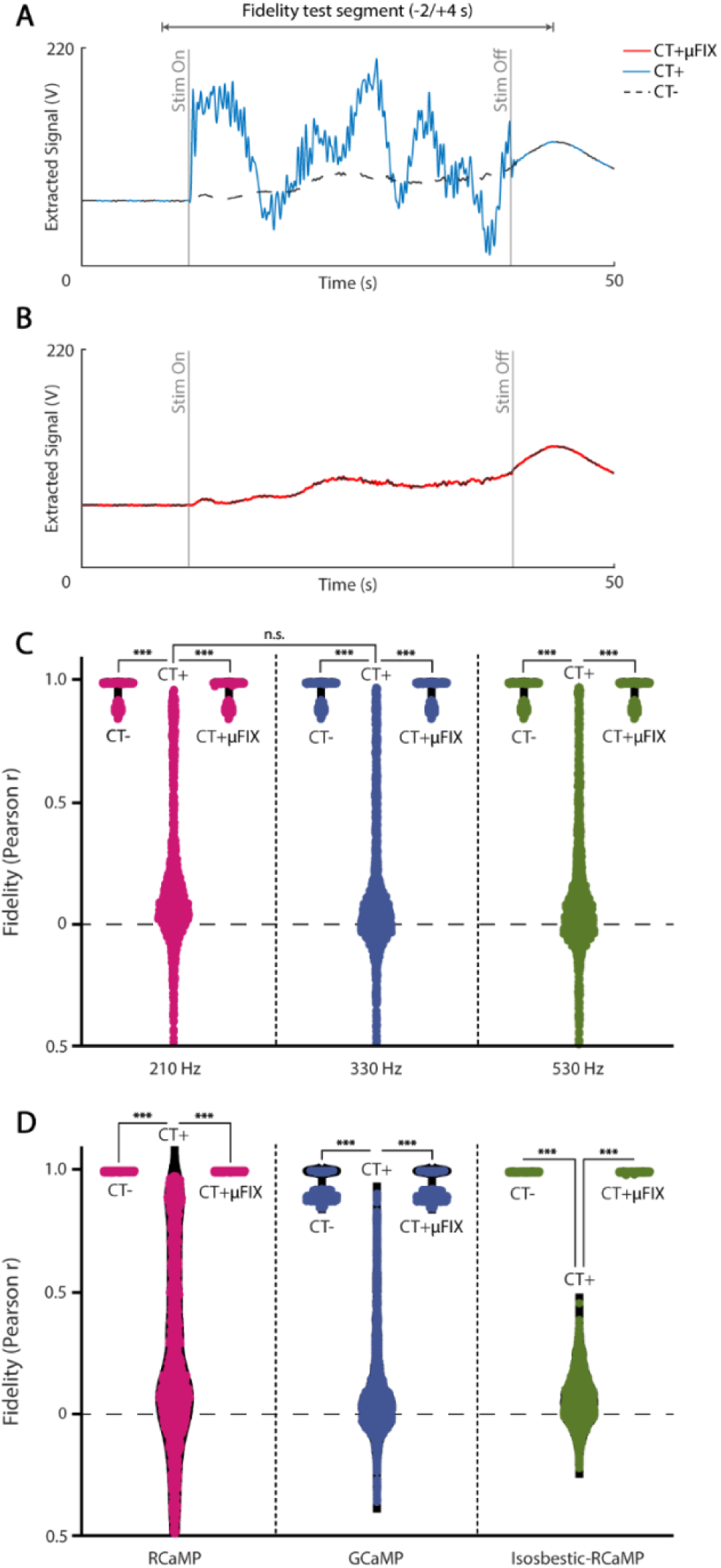
µFIX validation using simulated LIA with empirical responses as ground truth. **(A)** An example epoch of demodulated response CT+. **(B)** The demodulated response CT+µFIX for the same epoch as (C). **(C)** Pearson correlations between the ground truth and the LIA demodulated responses as described in Figure 4A. Results show simulation runs using 473 empirical recording epochs. We extracted up to three photometry responses from each epoch and ran the simulation three times with all responses rotated through the three carrier frequencies (210, 330 and 530 Hz). Altogether, there are 1605 matched data points for each group. **(D)** Grouping of data in (C) based on source channel, where RCaMP encodes the slowest-moving dynamics, GCaMP is faster with lower amplitudes, and Isosbestic-RCaMP encodes baseline noise without responses to stimulation.

**Table 2.**
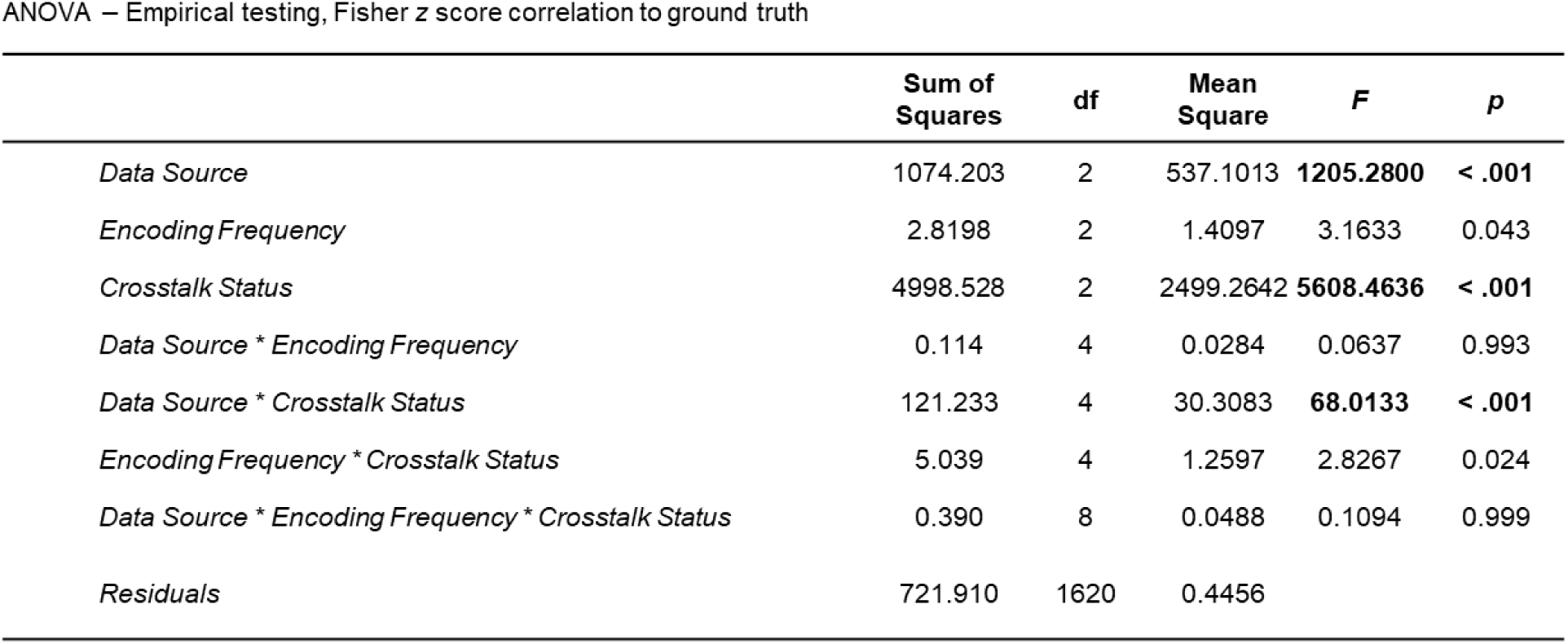
ANOVA for multi-variable influence on the recoverability of data from crosstalk. Data Source = Seizure response, flat response, or isosbestic data. Encoding Frequency = 210, 330, or 530 Hz encoding. Crosstalk Status = No crosstalk added (CT-), crosstalk added and not removed (CT+), or crosstalk added and removed (CT+µFIX).

The source of ground truth data significantly affected recovery fidelity (Table 2 ANOVA, *F* = 1205.28, *p* < 0.001). The signal source effect was most pronounced in CT+ results. Although the effect is still apparent on CT+µFIX in the Fisher Z-transformed fidelity scores, it translated to minor separation in group-wise medians (0.995 to 1.000). In particular, we note that signal recovery is practically equivalent regardless of the Ca2+ photometry variant (RCaMP or GCaMP), targeted neural population (excitatory or PV+), or whether the signal contained only a single active photometry source (RCa1) versus two simultaneous active sources (RCa2 and GCa2). Most of the effect between signal sources can be explained by a linear relationship between the fidelity score (Fisher z correlation coefficients) and the logarithm of the signal standard deviation (SD). Signals sourced from isosbestic channels typically had lower signal SD, which resulted in lower fidelity scores. For RCaMP and GCaMP sources, seizures often drive major changes in the source signals, leading to larger signal SD, resulting in high recovery fidelity. Based on our results, a signal SD of about 0.18 mV is required for signal restoration fidelity greater than 0.99 (Fig. S3).

While it is impossible to verify whether µFIX has accurately recovered the unknown physiological signaling values of empirical photometry from experimental recordings with real crosstalk, the present data strongly indicate that µFIX recovers data following similar simulated crosstalk. Further, our simulated results are likely generalizable since the result is identical when ground truth data are recovered from real or simulated crosstalk. Thus, we infer that, in most cases, µFIX can accurately recover photometry recordings when real optogenetic crosstalk is present.

### Computational cost and accuracy of µFIX compared to alternative approaches

Next, using our simulated LIA photometry pipeline and artificial ground truth, we compared µFIX with more uncomplicated crosstalk interpolation strategies to demonstrate its superior effectiveness in recovering the LIA photometry signal and assess its relative computational cost. We examined whether linear interpolation, spline interpolation, and single-frequency (1-Freq) interpolation are sufficient to recover the LIA photometry signal. Linear interpolation tests the approach of a straight line connecting across the ends of the saturated segment (Fig. 7D). Spline interpolation avoids demodulation artifacts arising from sharp transitions at the ends of the linearly interpolated segment (Fig. 7E). Lastly, we test if a single-frequency sinusoid using the lowest component carrier frequency (210 Hz) is sufficient to recover the photometry signal rather than the complete multi-frequency set (Fig. 7F).

**Figure 7.**
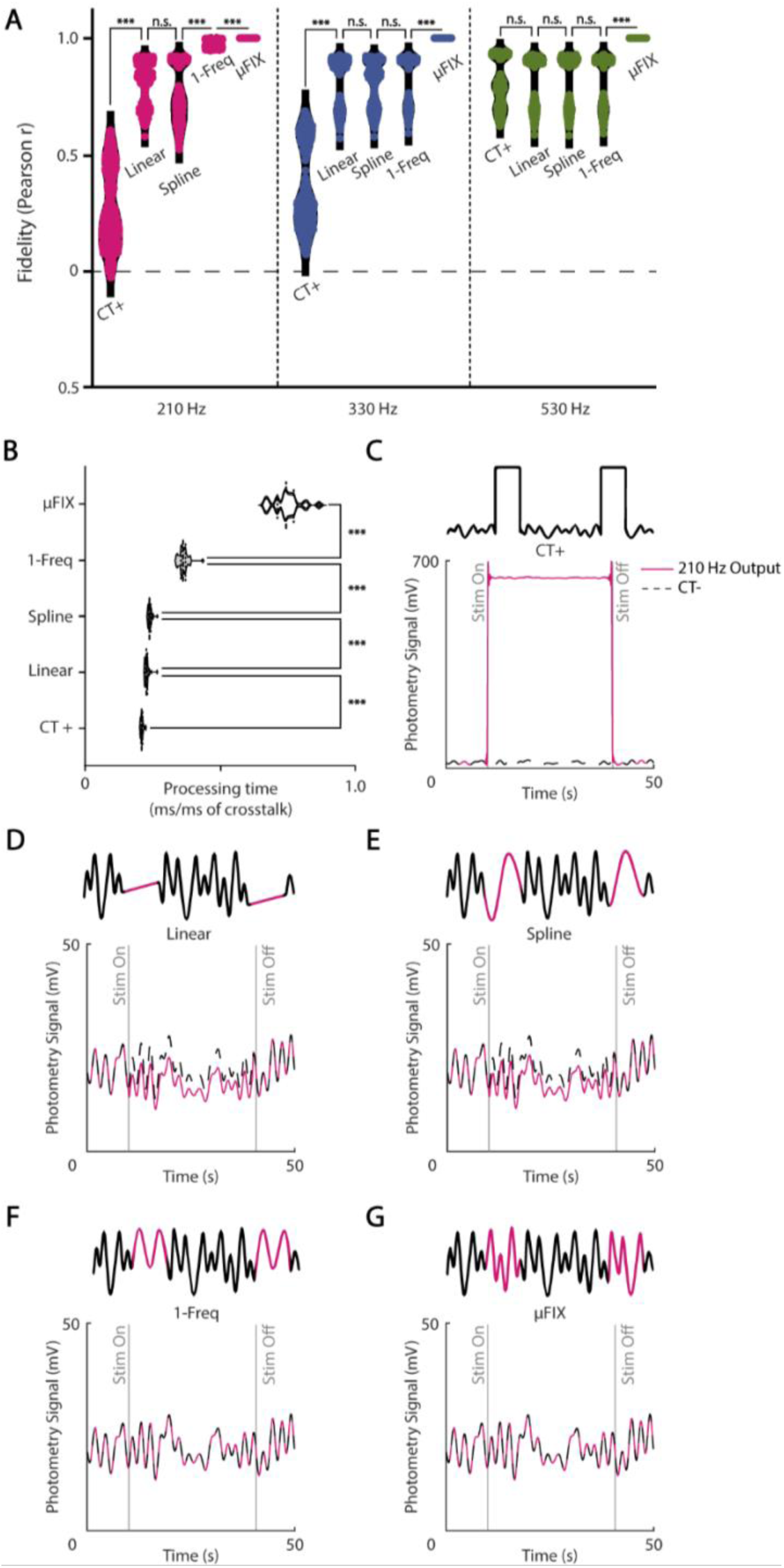
Alternative interpolation approaches for crosstalk snippet removal. **(A)** Pearson correlation by carrier frequency between the ground truth and the LIA demodulated response with simulated crosstalk (CT+), crosstalk segment interpolated with a straight-line (Linear), cubic Spline, single- frequency sinusoid (1-Freq), and µFIX. Results from simulation on the same artificial ground truths as in Figure 4. **(B)** Comparison of processing time for each interpolation approach expressed in milliseconds to process each cropped millisecond of crosstalk. **(C)** An example stimulation epoch of the demodulated response with simulated crosstalk (CT+). **(D)** The demodulated response after Linear interpolation for the same epoch as (C). **(E)** The demodulated response after Spline interpolation for the same epoch as (C). **(F)** The demodulated response after 1-Freq interpolation for the same epoch as (C). **(G)** The demodulated response after µFIX interpolation for the same epoch as (C).

Our results show that linear, spline, and 1-Freq approaches are ineffective in removing all the artifacts from crosstalk contamination. The LIA-demodulated response from the linear and spline interpolation illustrated a significant step change in the output but did succeed in removing the major unrecognizable segment of noise (Fig. 7D & 7E). This reflects the loss of signal at the carrier frequencies that were not replaced by these crosstalk interpolating methods. Overall, the fidelity of the demodulated output is 0.88 [0.64–0.92] for linear interpolation, 0.84 [0.61–0.92] for spline, and 0.91 [0.66–1.00] for 1-Freq interpolation; all were significantly lower than 1.00 [1.00–1.00] for µFIX (p < 0.001, ANOVA, Tukey’s post-hoc test).

Further, we wanted to see if the LIA encoding frequency of a signal makes simpler interpolation methods comparable to µFIX. Differences in fidelity score for µFIX in correlation units are indistinguishable (∼1.00), but the simpler approaches had significant variation. For 1-Freq interpolation, as expected, high recovery fidelity was found only for signals modulated on the fitted carrier frequency (210 Hz, 0.99 [0.95–1.00]), but not for signals on 330 Hz (0.90 [0.65–0.92]) or 530 Hz (0.87 [0.64–0.92]). For all other interpolation methods, signals on 530 Hz (0.90 [0.64–0.92]) and 330 Hz modulation (0.87 [0.64–0.92]) were restored much with higher fidelity than for 210 Hz (0.79 [0.59–0.95]) (p < 0.001, ANOVA, Tukey’s post-hoc test).

We compared the computational cost among these interpolation methods (Fig. 7B). On our virtual Windows server running on eight 18-core, 36-thread Intel Xeon 6254 processors @ 3.1 GHz, we estimated the time for each interpolation to process crosstalk segments from 20 simulated experiments. Each experiment was 30 min long and contained 10 epochs, each with 600x 5-ms crosstalk segments to process (20 Hz stimulation). Without applying any crosstalk recovery algorithm, each recording took 19.0 ± 0.4 (SD) ms to process each second of the recording. The µFIX algorithm was the most complex and took an average of 67.1 ± 4.5 ms/s, significantly higher than for simpler interpolation approaches (p < 0.001, ANOVA, Tukey’s post-hoc test). On average, µFIX took ∼30 s longer to process each recording; dividing this over the 6000 crosstalk segments, we estimate that each 9 ms crosstalk segment required ∼5 ms to process on our machine. This processing time may be short enough for deployment in the recording procedure so that Ca^2+^ photometry can be restored from crosstalk in real time.

### µFIX is accurately recovers signal from a wide range of stimulation protocols

The most effective optogenetic stimulation protocol varies depending on the opsin, target, and response of interest. This includes the duration of stimulation pulses, the frequency of the pulse train, and the length of the pulse train. So far, we have examined using 5 ms pulses at 20 Hz for 30 s of stimulation, which is optimal for generating seizures from the hippocampus. µFIX is designed to work with a wider range of stimulation protocols.

We sought to determine how well µFIX can recover the photometry response with longer durations of optogenetic stimulation. The quality of µFIX recovery was examined using our LIA simulation pipeline from 15 ms to 35 ms of crosstalk driven by a 20 Hz pulse train (Fig. 8A).

**Figure 8.**
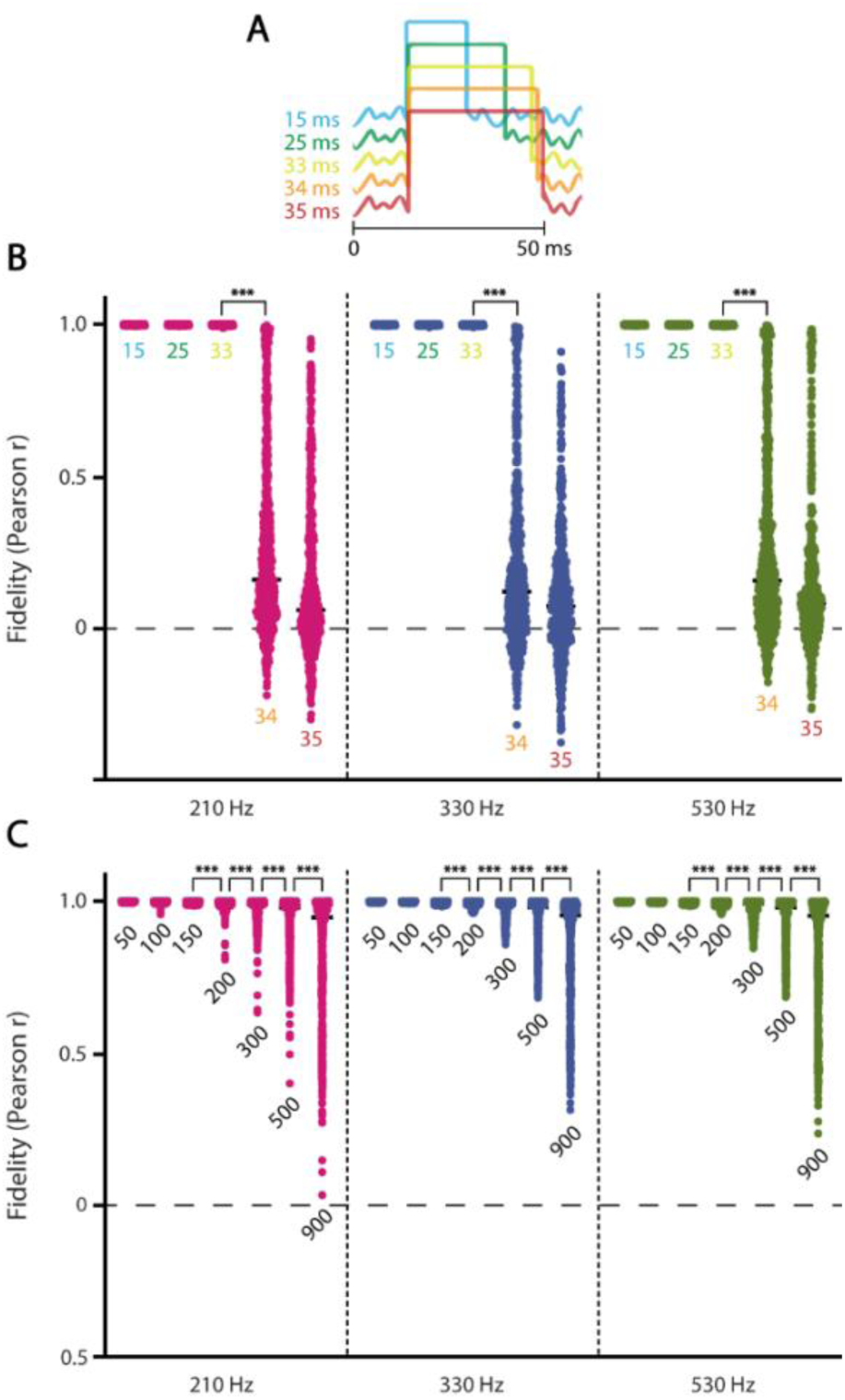
µFIX crosstalk recovery with longer pulse widths. **(A)** Illustration of stimulation pulse lengths corresponding to labels in panel (B). **(B)** Pearson correlations by carrier frequency between the demodulated response CT+µFIX (as described in Figure 4A) and the ground truth. Results from simulation on the same empirical-based ground truths as in Figure 5. Stimulation pulse frequency was at 20Hz (50ms pulse periods). Numeric labels for each group indicate the duration of the stimulation pulse in milliseconds. Insufficient of intact between pulses led to poor µFIX recover for pulses larger than 33 ms. **(C)** Same as (B), but with pulse frequency of 1 Hz and expanded pulse lengths. µFIX faithfully recovers stimulation pulses as large as 150 ms.

The fidelity of µFIX recovery was consistently maintained until a precipitous drop from 1.00 [0.99–1.00] with a crosstalk duration of 33 ms to 0.15 [-0.11–0.97] with a crosstalk duration of 34 ms (Fig. 7B, p < 0.001, ANOVA, Tukey’s post-hoc test). This is consistent with our algorithm’s requirement of ∼12 ms (2.5 cycles of 210 Hz) of intact recording preceding and succeeding the crosstalk segment for estimating the interpolation parameters. At 50 ms intervals between pulses (20 Hz), a 34 ms saturated segment with 4 ms of post-stimulation artifact is at the 12 ms limit, resulting in defective signal recovery.

Beyond the limit of intact data for 20 Hz stimulation, we sought the longest recoverable crosstalk segment by repeating the simulation with longer crosstalk durations using a 1 Hz pulse train (1000 ms intervals). We found that the average recovery fidelity reduced below 1.00 [0.97–1.00] for crosstalk durations longer than 200 ms, to 0.99 [0.88–1.00] at 300 ms, then gradually down to 0.95 [0.41–1.00] at 900 ms (Fig. 8C). Signals on the 210 Hz carrier frequency had significantly lower fidelity scores than 330 and 530 Hz (p < 0.001, ANOVA, Tukey’s post-hoc test). However, the reduction in recovery fidelity did not suffer an immediate drop compared to reaching the limit of the intact data (Fig. 8B). The sustained fidelity score with long crosstalk segments reflects the slow dynamics of the photometry response.

The highest frequency pulse train that µFIX can recover is limited by the length of intact recording in between pulses as the basis for a good estimate of the signal lost to crosstalk. As described, our standard algorithm requires 2.5 cycles of data at the lowest carrier frequency before and after each crosstalk segment for signal recovery, which is equivalent to ∼12 ms using a 210 Hz carrier. With 5 ms pulses, the highest pulse frequency under this setting is ∼50 Hz, already approaching the limit for the physiological firing rate of most neurons. Nonetheless, our simulations indicate that this is a very conservative setting. High-fidelity signal recovery can be achieved using as little as 1/4 cycles between pulses (∼1.2 ms at 210 Hz, fidelity = 1.00 [1.00– 1.00], Fig. S4), potentially working with pulse trains up to 100 Hz (with 5 ms pulses).

Lastly, as µFIX works on the level of an individual stimulation pulse and its induced crosstalk, we expect that the recovery fidelity would be independent of the length of the stimulation train. To verify this, we performed simulations with longer 5 ms, 20 Hz pulse trains lasting 60 s and 90 s (Fig. S5). As expected, we found no statistically significant contribution to recovery fidelity from the train duration main factor (p = 0.43, ANOVA).

## Discussion

The all-optical approach to interrogating neural populations enables cell-type-specific circuitry manipulation and activity read-out using genetic tools. Despite employing state-of-the-art optical spectra filtering (Fig. 1), optogenetic crosstalk contamination hampered our experimentation with multiple optical components, leading to distorted photometry recordings. Consequently, we developed µFIX to recover the underlying LIA fiber photometry signal from crosstalk. µFIX generates signal snippets of LIA-modulated fluorescence emission to replace noise-contaminated segments in the photosensor recording (Fig. 3). Perfect signal recovery fidelity (*r* ∼ 1.00) was achieved on non-crosstalk segments of a recording as well as simulated crosstalk-contaminated LIA photometry with ground truth signals of both artificial and empirical origins. Applying µFIX to our mice recordings effectively recovered photometry responses that resemble the dynamics of the same cell-type response when recorded from a reduced viral preparation that did not suffer from crosstalk.

### µFIX empowers experimental designs exposed to crosstalk

LIA photometry offers the benefits of multiplexing isosbestic reference signals^16,21,23–25^ and maximizes signal-to-noise ratio in fiber photometry^15,25,26^, but it is vulnerable to interruptions such as optogenetic crosstalk. In standard setups, a system of dichroic mirrors and advanced optical filters is employed to separate and route the emission spectra to different photosensor light paths (see our system in Fig. 1 as an example). These optical filters typically have a passband of 10-40 nm and an attenuation of OD5 (intensity reduction of five orders of magnitude) outside the passband. Higher OD and narrower passband filters can improve the filtering outcome but are likely to do so at the cost of some signal loss. However, light sources and fluorescent emissions typically have broad and possibly overlapping spectra, therefore, a complete signal separation by spectral filtering is not possible (see Fig. 3A). Using an optical filtering system is also insufficient to eliminate optogenetic crosstalk in the fiber photometry light path when the employed opsin and fluorescence indicator have overlapping spectra (e.g., ChRmine and RCaMP).

In our recordings, in spite of optical filtering and spatial separation of the fiber photometry cannula from the optogenetic stimulation cannula, optogenetic crosstalk was still evident. It continued to interrupt the carrier signal in square-wave patterns, resulting in characteristic artifacts in the demodulated photometry response. The sharp signal transitions into and out of the stimulation periods produced a large, transient, and ripple-like “ringing artifact” at these time points. During the stimulation period, depending on whether the content at each carrier frequency was boosted or masked out, crosstalk also caused a step-like change in the mean response level. The frequency of stimulation also induced a beat frequency oscillation that can overwhelm the photometry response (Fig. 2A & B). The effect of crosstalk on the demodulated response ranged from severe to subtle, depending on the interaction between the stimulation frequency, LIA carrier frequency, and whether crosstalk saturates the photosensor. Recognizing even the subtle artifacts is imperative, as they can be mistakenly interpreted as stimulus-driven responses.

Whether or not optogenetic crosstalk is present, µFIX is a convenient tool that can be applied indiscriminately to any suspected LIA photometry recording without producing adverse effects, providing there is sufficient response amplitude driving the fluorescence signal (0.3 mV standard deviation, Supplemental Fig. 1). µFIX does not distort a response absent of crosstalk (Fig. 2 & 3). Where crosstalk is present, we demonstrated that µFIX achieves perfect recovery (r ∼ 1.00) for a wide range of stimulation durations (up to 150 ms long, Fig. 7) and signals with slow or fast dynamics (up to 20 Hz, Fig. 5 and S2). To our knowledge, this encompasses the majority of experimental designs using optogenetics and fiber photometry. With a processing time of < 1 ms per ms of crosstalk noise (Fig. 6B), µFIX may be deployed in the real-time recording pipeline during experimentation.

Fiber photometry is typically only capable of resolving slowly changing bulk signaling from the intended targets^5,6,8,27^ well within the 20 Hz working limit of the µFIX algorithm. Our empirical signals of interest have the optimal signal-to-noise ratio with a 3 Hz low-pass filter. In a similar application using widefield imaging in monkey V1^28^, the visually-driven response in GCaMP6 diminished sensitivity above 4 Hz stimulation. In two-photon imaging, calcium indicators for capturing single-neuron activity are typically recorded with a frame rate of less than 40 Hz^29,30^, i.e. dynamics up to 20 Hz. The working limits of µFIX encompass with the maximum kinetics of the current, commonly used indicators.

Adopting µFIX requires short optogenetic stimulation pulses. A non-exhaustive survey within our research interest (seizures and epilepsy) indicated that most researchers are adopting stimulation pulses between 5 to 20 ms ^9,14,31–34^. Short optogenetic stimulation pulses have also proven successful at eliciting neural activity and are a widely adopted stimulation technique^35,36^. The effective stimulation pulse duration depends on the specific protocol and opsin, with up to 2000 ms reported ^37^. For stimulation protocols that utilize constant light delivery, such as those typically for activating inhibitory networks, only minimal modification is required to create an equally effective high-duty-cycle pulse-train paradigm compatible with µFIX^38,39^.

As a solution to optogenetic crosstalk, µFIX makes it possible to design experiments with fluorescence photometry sensors whose spectra overlap the spectrum of optogenetic stimulation light. Investigators can simultaneously utilize the blue and red-spectrum fluorescence channels (e.g., GCaMP and RCaMP) to monitor the fiber photometry responses of two neural populations or neurotransmitters under optogenetics manipulation. An example of a design with dual photometry and optogenetics is demonstrated in our experimental setup (Fig. 1), which was produced to study the relative excitatory/inhibitory signaling balance leading to a perturbed state of the hippocampus that produces seizures (Fig. 2). In addition, with the use of µFIX, investigators are unrestricted from employing the most suitable optogenetic opsin for their experiment, including the combination of blue and red-spectrum opsins for dual-optogenetic manipulation control (e.g. both ChR2 excitation and NpHR inhibition) alongside simultaneous fiber photometry.

µFIX may also combat signal interference wherever LIA (or optical lock-in detection) is used, including other imaging methodologies. Some of the potential use cases are voltage-sensitive-dye imaging^40^, fluorescence-detected multidimensional electronic spectroscopy^41^, infrared microscopy^42^, Raman microscopy^43^, and immunofluorescence microscopy^44^.

### Limitations and future extensions of µFIX

High-fidelity signal recovery with µFIX is built on the fundamental mathematic principles behind multi-frequency LIA photometry. Lesser interpolation approaches (linear, spline, or single-frequency interpolation) did not produce the same fidelity levels (Fig. 7). While µFIX is robust in its current form (see Fig. 5, 8, S4, S5), and more than sufficient for the majority of applications that we are aware of, assumptions were made in the design of the µFIX algorithm may pose constraints on experimental designs using indicators with extremely fast kinetics or uncommon optogenetic pulse train patterns.

First, to simplify the µFIX formulation, we assumed that the amplitude of the modulated signal is constant during the interpolated segment. This assumption was made because the dynamics of the signal of interest from fiber photometry recordings are generally slow (subsecond scale), and the crosstalk segment is short (milliseconds). While this assumption simplifies µFIX implementation, it has implications for removing crosstalk from stimulation protocols requiring > 150 ms pulse duration (maybe working with inhibitory opsins) or restoring signals with very fast > 20 Hz dynamics (such as voltage-sensitive indicators). For applications working beyond these limits, we believe extending µFIX to model a dynamic amplitude signal profile, such as a cubic spline function or inspiration from deep learning image inpainting approaches^45^, will allow recovery for more extended pulse widths and faster signal dynamics.

Secondly, µFIX requires uncontaminated data to fit its model parameters for reconstructing the signal to crosstalk. We used a conservative 2.5 cycles of the lowest carrier frequency as the required duration of uncontaminated data before and after each contaminated snippet. Our simulation indicates that as little as 1/4 cycles may be used without compromising crosstalk signal recovery (Fig. S4). This imposed a minimum off time between stimulus pulses of ∼1.2 ms on the 210 Hz carrier and subsequently limits the duty cycle of the stimulation, a consideration for adapting experiments with constant light delivery to use µFIX. This will also impact experiments requiring a stimulus frequency of >100 Hz. The minimum off time can possibly be reduced by adopting higher LIA carrier frequencies – e.g. with a 530 Hz carrier, which reduces it to 0.47 ms, but we have not tested this setting.

### Informing the choice of LIA carrier frequency for photometry

Carrier frequencies typically used for LIA fiber photometry (and provided as defaults by equipment vendors such as TDT Systems) are 210, 330, and 530 Hz (Fig. 3A). The frequency choices are relatively narrow; they must avoid line noise (50 or 60 Hz) and its harmonics as well as not interact with other carriers and their harmonics. In all our simulations, we multiplexed and re-extracted three different signals respectively modulated at 210, 330, and 530 Hz, and our results confirmed that these are a suitable mixture of carrier frequencies for LIA photometry. We did not encounter any interference between these carrier frequencies (although we did not specifically test for this). In general, signals were encoded/decoded equally well on all three frequencies (Figs. 4B, 5B, 6C, 8B, 8C). There is evidence that 530 Hz may be better at encoding faster signal dynamics (up to 100 nps or ∼50 Hz, Fig. 5), which is generally not required for capturing bulk fluorescence activity from calcium and GRAB sensors but may be helpful in specific experimental scenarios^46^. While it remained essential to choose the appropriate carrier frequencies in the experimental setup, our results indicated these choices had little impact on encoding quality, immunity from crosstalk, and the performance of µFIX.

Spectral analysis of in vivo recordings employing 210, 330, and 530 Hz carrier frequencies indicate that there may be non-linear interactions between them. Spectral peaks other than the carrier frequencies were present (Fig. 3A). Some were observed at the harmonics of the carrier frequency (420Hz and 660Hz). We suspect this originated from the non-linear excitation-emission response of our chosen calcium indicators. However, an alternate explanation is an imperfect sinusoidal modulation of the excitation light or non-linear loss in light transmission. We also observed a spectral peak at 120 Hz. We believe this was not a harmonic of the line noise (no corresponding 60 Hz peak); instead, it was the beat frequency between the 210 and 330 Hz carrier frequencies. However, the beat frequency is a time domain manifestation between two sinusoidal signals and does not have a spectral peak; one at the beat frequency indicated a non-linear mixing of the two fluorescence emissions, either during transmission or at the Ca^2+^ indicator. The actual source of these spectral peaks requires further investigation.

### Alternative fiber photometry paradigms to avoid optogenetic crosstalk

Time-division multiplexing (TDM) photometry is an alternative to LIA photometry. TDM involves alternating frames of each component light source in the experimental setup so that only one fluorescence is excited and recorded at a time^16,47–49^. TDM can partition optogenetic stimulation in separate time slots to avoid crosstalk. Although there is evidence that TDM may be more noise-tolerant than LIA photometry^16^, a more rigorous comparison between these two techniques is required to confirm its advantage. The major disadvantage of TDM is that it requires stimulation to be within pre-determined time divisions, which may hinder the flexibility of the stimulation paradigm. Finally, TDM requires additional specialized hardware and software for researchers who have already adopted LIA photometry systems.

Fiber photometry can also be conducted using a constant-level excitation light, i.e., without LIA or TDM modulations. Brief, crosstalk-affected segments at the photosensor can be bridged with simple linear or spline interpolation. However, continuous excitation photometry does not benefit from the improved signal-to-noise ratio of LIA encoding and would suffer greater noise throughout the entire recording. Additionally, this setup would not allow a simultaneous isosbestic reference to be recorded, which is required to correct fluorescence deviations of non-neural origins in the photometry response.

## Conclusion

We developed µFIX and showed that it is an effective method to recover LIA-encoded photometry signals contaminated by optogenetic crosstalk. We demonstrated that µFIX allows a robust optogenetics stimulation paradigm and is computationally efficient for real-time implementation. µFIX enables extended experimental designs employing multiple simultaneous fiber photometry and optogenetics channels to study the neural circuitry in a previously unfeasible manner due to crosstalk.

## Methods

### Animal Preparation

Experiments were carried out with adult (20-25 weeks old) male and female inbred homozygous PV-Cre mice (B6.129P2-Pvalb<tm1(cre)Arbr>/J – The Jackson Laboratory, Bar Harbor, ME, USA). Mice were reared under social housing and environmental enrichment conditions with ad libitum food and water, standardized 12 h light/12 h dark cycle (lights on at 06:00 AM), temperature, and humidity. All animal studies were conducted via Rutgers IACUC within an AAALAC-accredited facility. IACUC Protocol #PROTO201900218.

Data from four animals were used (Table 1). All animals underwent stereotaxic surgeries to receive intrahippocampal adeno-associated viral vector (AAV) injections and a headcap optrode ensemble implantation. AAVs targetting the excitatory pyramidal cells were microinjected in the left dorsal hippocampal CA1 (AP -2.1; ML -1.6; DV -1.4) for optogenetic stimulation. In the right dorsal hippocampal CA1 (AP 2.1; ML 1.6; DV -1.4), a single AAV (mice OP191 & OP193) or an AAV cocktail for dual photometry (OP2718 & OP275) was injected to read out the Ca^2+^ activity of PV interneurons and the excitatory pyramidal cells. All coordinates are given in millimeters from bregma: anterioposterial (AP), mediolateral (ML), dorsoventral (DV).

The exact viral vector combination used varied between mice. Two of the animals, OP275 and OP2718, received 300nL of bicistronic ChRmine (AAV-8-CaMKIIa-GCaMP6m-p2a-ChRmine- TS-Kv2.1-HA; GVVC-AAV-180, Gene Vector and Virus Core, Stanford, CA, USA) on the left hippocampus, with 300 nL of Cre-dependent GCaMP6 (pAAV.Syn.Flex.GCaMP6m.WPRE.SV40; #100838, Addgene, Watertown, MA, USA) and 100 nL of RCaMP (AAV8-CaMKIIa-JRCaMP1b; GVVC-AAV-150, Gene Vector and Virus Core) in the right hippocampus. The biscistronic ChRmine vector transfects the same neurons with GCaMP for Ca^2+^ activity readout. OP191 and OP193 received 300 nL of ChR2 (pAAV-CaMKIIa-hChR2(H134R)-EYFP (AAV5); #26969, Addgene) in the left hippocampus with only a single GECI in the right hippocampus. OP193 was targeted for contralateral PV interneurons using Cre-dependent RCaMP (pAAV1.Syn.Flex.NES-jRCaMP1b.WPRE.SV40; #100850, Addgene). OP191 was targeted for contralateral CaMKII cells using RCaMP (AAV8-CaMKIIa- JRCaMP1b). Injections were made at a rate of 30 nL/min; the needle was maintained in place for 5 minutes after injection to avoid backflow and then slowly retracted.

Animals were implanted with a headcap optrode ensemble (Fig. 1C) either in the same surgery (OP191 & OP193) or in a second surgery (2 – 13 weeks after, see Table 1). The head cap ensemble was composed of the two optic fiber cannula (0.66 numerical aperture, 400 um diameter, Doric Lenses, Québec, QC, Canada) sitting just above the bilateral viral vector injection site (AP ±2.1; ML -1.6; DV -1.1), and colocated CA1 (DV -1.4) and DG (DV -1.9mm) recording electrodes (4 altogether, EM6/3/SPC, P1 Technologies, Roanoke, VA, USA).

Electrodes were interfaced to a 6-channel connector pedestal (mini6 format, P1 Technologies) with two additional reference electrodes (EM6/96/1.6/SPC or EM6/3/SPC, P1 Technologies) above the prefrontal cortex (AP 1.6; ML ±1.6) or the cerebellum (AP -5.0; ML ±1.6). The ensemble was secured to the skull using dental acrylic cement.

Animals were maintained on isoflurane anesthesia (∼1.5% isoflurane in pure oxygen) on a heating pad during the surgeries. Sustained-release Buprenorphine (Ethiqa XR, 3.25 mg/kg, s.c.; Ethiqa XR, North Brunswick, NJ, USA) was administered to the animals to alleviate pain and discomfort after recovering from the procedure. After the surgeries, animals were transferred to a clean cage to recover and single-housed.

### Perfusion and Histology

After experimentation, the animals were deeply anesthetized with isoflurane and perfused transcardially with 4% paraformaldehyde (PFA) in 1X phosphate buffer (PBS – pH7.4). Brains were extracted and post-fixed for at least 24 hours in 4% PFA and then transferred to 30% sucrose with 0.02% sodium azide in 4% PFA for at least two days at 4°C. Coronal brain sections of 30μm thickness were made on a cryostat (Leica CM3050S, Leica Microsystems, Wetzlar, Germany). Sections were mounted on slides and cover-slipped with DAPI mounting medium.

Fluorescent images of the viral expression were acquired on a Leica fluorescent microscope (Leica DM 4B; Leica Microsystems, Wetzlar, Germany).

### Animal Recording

Recordings were performed at least 22 days after the injection surgery to allow time for viral expression. Simultaneous electrophysiological recording, optogenetic stimulation, and Ca^2+^ fiber photometry were conducted using a TDT system (RZ10x, PZ5, RA16; Tucker Davis Technologies, Alachua, FL, USA). ChRmine optogenetic stimulation was delivered using either the inbuilt RZ10x 590 nm LED (Lx590) or a 589 nm laser (LMS-BY02-GF3-00020-05, Laserglow Technologies, Toronto, ON, Canada). ChR2 optogenetic stimulation was delivered using the inbuilt RZ10x 465 nm LED (Lx465). Excitation light for fiber photometry was delivered with inbuilt RZ10x LEDs: 405 nm for isosbestic signal (Lx405), 465 nm for GCaMP (Lx465), 560 nm for RCaMP (Lx560). 6-port mini cubes (FMC6_IE(400-410)_E1(460- 490)_F1(500-540)_E2(555-570)_F2(580-680)_S, Doric Lenses) was used to merge and split the light paths (Fig. 1).

For mice OP2718 and OP275, we interfaced the ChRmine optogenetic light and excitation lights to the left fiber cannula for isosbestic and GCaMP signals. Optogenetic stimulation was delivered through the RCaMP emission port (F2) of the Doric mini cube. We interfaced the excitation lights to the right fiber cannula for isosbestic, GCaMP, and RCaMP signals. The returning emissions from the animal were optically split by the GCaMP and RCaMP spectra and routed to separate photosensors on the TDT system. For OP191 and OP193, the right fiber cannula was similarly interfaced for photometry, while the ChR2 optogenetic light was directly routed to the left fiber cannula without the Doric mini cube. Both electrophysiological and photometry signals were recorded at 6.1 kHz.

Optogenetic stimulation was delivered to the left hippocampus to induce seizures in the animals. Each mouse was subjected to an extensive battery of stimulation experiments. This paper includes only experiments using 30-second stimulation trains with 10 Hz or 20 Hz 5 ms pulses. We refer to each 30 s stimulation episode as an epoch. In each experiment, multiple stimulation epochs were delivered, each separated by 90 s non-stimulation time (i.e., 120 s between the start of each epoch). The stimulation power was 2-10 mW, depending on the required power to induce seizures for each mouse in each experiment. This power was measured at the fiber optic tip with a power meter (PM20A, ThorLabs).

We collected 45 experiment recordings across four mice (see Table 1). These experiments have at least one seizure response and a demonstrated photometry response on either contralateral channel. Altogether, 535 stimulation epochs were collected, of which 206 resulted in seizures, and 329 did not.

### Lock-in Amplification (LIA) Photometry: Encoding, Multiplexing and Demodulation

LIA has been adopted in many fiber photometry applications. It is a signal-processing method that encodes the original signal (*V*_*s*_) with a carrier wave of a higher frequency (*f*_1_) to increase tolerance to noise in the course of signal transmission^16,26^:

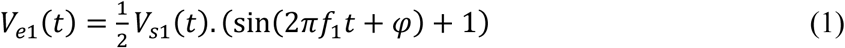

A notable strength of the LIA approach is the ability to multiplex multiple signals into a single transmitted signal (*V*_*T*_). LIA photometry exploits this LIA strength to encode the isosbestic reference^21,24^ and emission of multiple fluorescence indicators into the same fiber optic channel^26^. The multiplexing is achieved by encoding all component signals in orthogonal sinusoidal frequencies (Fig.1):

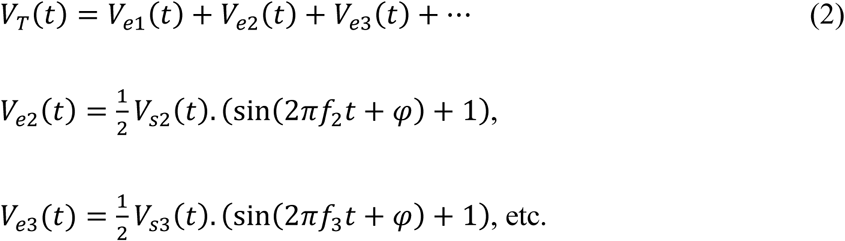

Leveraging the orthogonality of sinusoidal waves, the encoded signals (*V*_*e*_) can be extracted by isolating the transmitted signal (*V*_*T*_) at the desired encoding frequencies (*f*_1_, *f*_2_, etc.) using Euler’s formula.

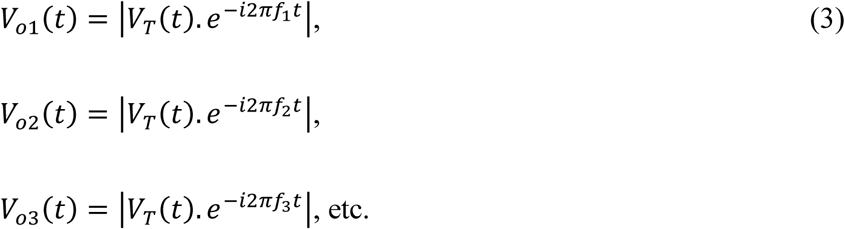

Lastly, the original signal (*V*_*s*_) can be demodulated from the extracted signals (*V*_*o*_) by averaging the signal over a sufficiently long time span to remove the carrier frequency. We implemented this using a fifth-order Butterworth low-pass filter with a cut-off frequency of 3 Hz.

In our animal experiment, we multiplexed up to three light excitation sources to a single optic fiber to read out both GCaMP (470 nm), RCaMP (560 nm), and their respective isosbestic signal (405 nm). Each light source was modulated with a different LIA carrier frequency from 210 Hz, 330 Hz, 450 Hz, or 530 Hz. The exact combination varied slightly between experiments. Our most common configuration was 330 Hz and 530 Hz for ipsilateral GCaMP and isosbestic, with 210 Hz, 330 Hz, and 530 Hz for contralateral GCaMP, RCaMP, and isosbestic, respectively.

Note that isosbestic references for GCaMP and RCaMP were excited with the same light source (and, by extension, carrier frequency). Their respective isosbestic signals were differentiated by the separate optic filter path for GCaMP and RCaMP and thus from the signal recorded from the separate photosensors.

### µFIX - Optogenetic Crosstalk Filling-in

µFIX (**Mu**lti-**F**requency **I**nterpolation **X**-talk removal algorithm) seeks to estimate the signal displaced by crosstalk (*V*^_*T*_) using the multi-frequency LIA transmission model described in Eq (2):

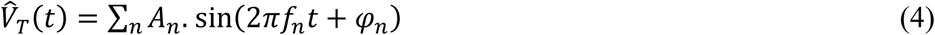

The amplitude (*A*_*n*_) and phase (*φ*_*n*_) of each carrier sinusoid (*f*_*n*_) were estimated from the intact signals just before and after each contaminated segment. We assumed a static *A*_*n*_ throughout each short segment filled in. Standard LIA demodulation was applied to the adjusted transmission signal to recover the photometry response.

For our application, we fitted the model to data representing 2.5 cycles of the lowest carrier frequency (210 Hz) before and after the contaminated segment (i.e., 12 ms or 72 samples at 6.1 kHz of data at each end). Model fitting was solved using Matlab’s nonlinear, least-squares algorithm (R2023a, MathWorks, Natick, MA, USA).

#### Alternative Interpolation Approaches for Crosstalk Recovery

We investigated simpler alternative approaches for filling in crosstalk segments. (1) A line connecting the ends of the contaminated segment; (2) a smoothed spline interpolating between the ends of the contaminated segment (using Matlab’s inbuild fillmissing() function), and (3) a single-frequency sinusoidal (1- Freq) interpolation at the lowest carrier frequency (210 Hz in our experiment). Single-frequency model fitting using the least-squares method in Matlab. These approaches were compared against µFIX for response recovery fidelity and computational time using the simulated LIA photometry pipeline with artificial ground truth. Testing was performed on Matlab R2023a running in Windows Server 2019 Standard (Microsoft, Albuquerque, NM, USA) as a virtual server with eight Intel Xeon Gold 6254 processors (18 cores, 36 threads, 3.1 GHz; Intel, Santa Clara, CA, USA) and 128 Gb of RAM. The time to process all interpolations and demodulation for the entire stimulus train (20 Hz of 5 ms pulses over 30 s, 600 segments altogether) was measured using Matlab’s inbuilt tic/toc functions.

### Simulated LIA Photometry Pipeline and Artificial Ground-Truth Signal

A simulation of the LIA photometry encoding process was devised using Eq (2). The pipeline is illustrated in Fig. 4A. The pipeline starts with the ground truth photometry response and simulates the LIA-encoded fluorescence read-out of the ground truth. Both artificially generated and empirical signals were inputs (ground truth) to the simulation pipeline.

For artificial ground truth, slow-varying signals were generated starting with 3 random numbers per second (nodes per second, nps) between 0 and 0.1 V. The signal amplitude and nps value were selected to approximate our empirical data. We retested with ground truth signals up to 150 nps to simulate faster signaling dynamics.

Each sequence of nodes was then interpolated to a smooth 1.0 kHz sampling rate signal (*V*_*s*_). Three independent artificial signals (triplet) were generated for each simulation run and up- sampled to 6.1 kHz for LIA encoding according to Eq (1) using carrier frequencies 210 Hz, 330 Hz, and 530 Hz, respectively, to match our empirical setup. The three LIA-encoded signals were then multiplexed into a single signal (*V*_*T*_) as described in Eq (2). LIA demodulation was performed on the multiplexed signal and then down-sampled to 1.0 kHz (*V*_*o*_) to match the 1.0 kHz ground truth (*V*_*s*_). These sampling rates were chosen to match our empirical recording hardware setup. To assess the effect of the carrier frequency, the modulation/demodulation test pipeline for each artificial ground truth triplet was repeated three times, with each signal assigned to a different carrier frequency in each repeat.

#### Simulating Optogenetic Crosstalk

Crosstalk was simulated by setting the carrier wave equal to 10 V for the duration of stimulation—this is the photosensor saturation limit. Real crosstalk exhibits a brief 2 ms post-stimulation rebound up to -1 V, which we also simulated. Crosstalk was inserted into the multiplexed signal (*V*_*T*_) in our simulation pipeline. To match our empirical optogenetic stimulation delivery, segments corresponding to a 30 s train of 5 ms pulses at 20 Hz were saturated to construct a crosstalk contaminated signal (*V*_*XT*_). We refer to the LIA demodulated response from the uncontaminated multiplexed signal as CT-, and the demodulated response from simulated crosstalk as CT+. µFIX was applied to the contaminated signal *V*_*XT*_ .

The subsequent demodulated response was named CT+µFIX. The same pipeline was used to quantify the effectiveness of photometry signal recovery using alternative crosstalk interpolation approaches (linear, spline, 1-Freq).

#### Measurement of Signal Recovery Fidelity

Output from LIA demodulation was compared using Pearson’s correlation coefficient. This is used to assess the severity of crosstalk-mediated signal distortion and the fidelity of the µFIX recovery. The correlation coefficient was calculated between two demodulated signals from 2 s before each 30 s stimulation train to 4 s after the stimulation (36622 samples at 6.1 kHz), during which we expect the crosstalk to affect their output. Due to its non-normal distribution nature, fidelity scores are reported using the median and [2.5–97.5]% quantile range in correlation units unless otherwise labeled. Statistical calculations were conducted on the Fisher Z-transformed value (*z*) of the correlation coefficients (*r*) to improve the normality of the data distribution:

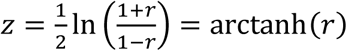

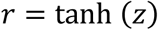

### Empirical Ground Truth Data Pool and Testing

We extracted epochs from our empirical recordings to form a pool of empirical ground truth data for the simulated LIA photometry pipeline. We picked three photometry channels from each mouse to form our empirical ground truth triplet in place of the artificial ground truth; otherwise, the same simulation and testing pipeline was used. We include the contralateral GCaMP, RCaMP, and the isosbestic GCaMP responses. We applied µFIX on all signals to obtain non- crosstalk contaminated photometry signals as the ground truth. Altogether, from 43 recordings across eight animals, we collected 208 epochs of photometry response triplets.

The fidelity of µFIX was validated by drawing on this empirical data pool as ground truths for the simulated LIA photometry pipeline. Testing was executed the same way as described for artificial ground truth, here substituted with three photometry responses from each empirical epoch: RCaMP, GCaMP, and GCaMP isosbestic. To isolate differences between carrier frequencies, we repeated the stimulation three times with the carrier frequency assignments rotated through the photometry responses, such that each photometry response was tested once at each of the three carrier frequencies 210, 330, and 530 Hz. This creates 1605 total empirical trials of µFIX.

#### Simulating Longer Duration Crosstalk Segments

The empirical data pool was used as the ground truth signal to assess the fidelity of µFIX recovery with longer crosstalk durations.

Simulations were conducted as previously described for empirical ground truth data. In the first batch of simulations, we varied the stimulation pulse duration from 5 to 35 ms using 20 Hz pulse trains. In the second batch of simulations, we reduced the pulse train frequency to 1 Hz to explore longer pulse durations up to 900 ms.

## Statistical Analysis

All Pearson r correlations (fidelity) statistics were conducted using Fisher-transformed values and inverse transformed for plotting. ANOVA was used to evaluate the contribution of data source, channel, encoding frequency, interpolation method, and crosstalk length to fidelity scores. ANOVA and post-hoc tests were performed in Jamovi (v2.4.14.0) using Tukey’s correction for multiple comparisons. Student’s paired t-test was used to compare the difference between means for processing time in our interpolation method pipeline. T-tests and violin plots were created using GraphPad Prism 10.

## Data and Source Code Availability

Code and documentation available at: https://github.com/maxbrkstone/mufix.

## Supporting information

PDF: Additional example recordings from two mice of before and after crosstalk removal via µFIX (S1), power spectral analysis of empirical and simulated epochs (S2), signal recovery fidelity vs. source signal standard deviation (S3), µFIX crosstalk removal results with varying length of intact data surrounding crosstalk (S4), µFIX crosstalk removal results in 60 and 90 second stimulation compared to the 30 second default (S5).

## Author Contributions

Conceptualization: S.C.C., H.S. Method: M.B., S.C.C. Software: M.B., S.C.C. Data curation: M.B., S.C.C., S.V., E.C., L.S.P., F.T. Investigation: M.B., S.C.C., S.V., E.C., F.T. Validation:

M.B., S.C.C. Formal analysis: M.B., S.C.C. Supervision: D.J.B., H.S. Funding: R.E.G., H.S.

Visualization: M.B., S.C.C., L.S.P., F.T. Project admin: S.C.C., R.E.G., H.S. Writing: M.B.,

S.C.C. Review: M.B., S.C.C., D.J.B., H.S.

## Acknowledgments

none

## For Table of Contents Only

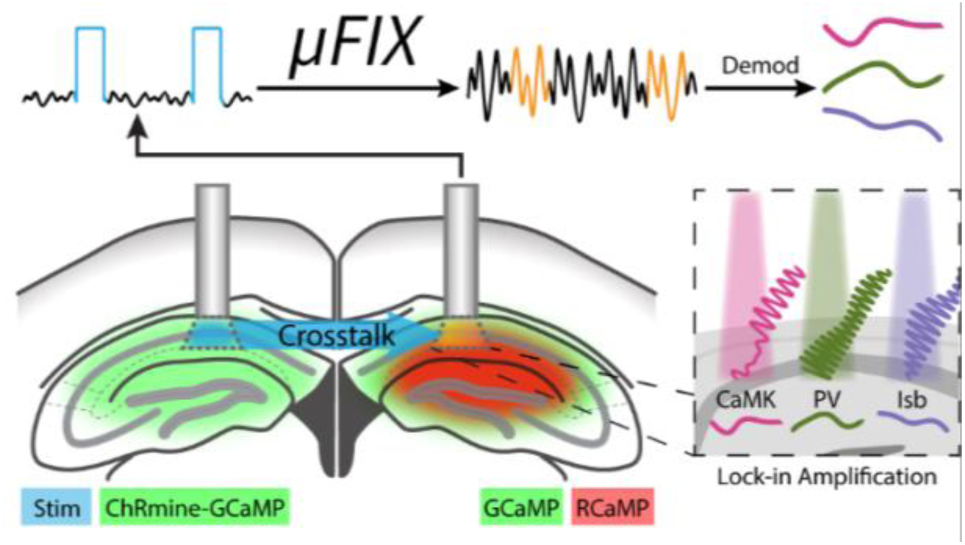

## References

(1) Zeng, H.; Madisen, L. Mouse Transgenic Approaches in Optogenetics. Prog. brain Res. 2012, 196, 193–213. 10.1016/b978-0-444-59426-6.00010-0.

(2) Sidor, M. M.; Davidson, T. J.; Tye, K. M.; Warden, M. R.; Diesseroth, K.; McClung, C. A. In Vivo Optogenetic Stimulation of the Rodent Central Nervous System. J. Vis. Exp. : JoVE 2015, No. 95, 51483. 10.3791/51483.

(3) Chen, W.; Li, C.; Liang, W.; Li, Y.; Zou, Z.; Xie, Y.; Liao, Y.; Yu, L.; Lin, Q.; Huang, M.; Li, Z.; Zhu, X. The Roles of Optogenetics and Technology in Neurobiology: A Review. Front. Aging Neurosci. 2022, 14, 867863. 10.3389/fnagi.2022.867863.

(4) Wick, Z. C.; Krook-Magnuson, E. Specificity, Versatility, and Continual Development: The Power of Optogenetics for Epilepsy Research. Front. Cell. Neurosci. 2018, 12, 151. 10.3389/fncel.2018.00151.

(5) Wang, Y.; DeMarco, E. M.; Witzel, L. S.; Keighron, J. D. A Selected Review of Recent Advances in the Study of Neuronal Circuits Using Fiber Photometry. Pharmacol. Biochem. Behav. 2021, 201, 173113. 10.1016/j.pbb.2021.173113.

(6) Simpson, E. H.; Akam, T.; Patriarchi, T.; Blanco-Pozo, M.; Burgeno, L. M.; Mohebi, A.; Cragg, S. J.; Walton, M. E. Lights, Fiber, Action! A Primer on in Vivo Fiber Photometry. Neuron 2024, 112 (5), 718–739. 10.1016/j.neuron.2023.11.016.

(7) Inoue, M. Genetically Encoded Calcium Indicators to Probe Complex Brain Circuit Dynamics in Vivo. Neurosci. Res. 2021, 169, 2–8. 10.1016/j.neures.2020.05.013.

(8) Wu, Z.; Lin, D.; Li, Y. Pushing the Frontiers: Tools for Monitoring Neurotransmitters and Neuromodulators. Nat. Rev. Neurosci. 2022, 23 (5), 257–274. 10.1038/s41583-022-00577-6.

(9) Khoshkhoo, S.; Vogt, D.; Sohal, V. S. Dynamic, Cell-Type-Specific Roles for GABAergic Interneurons in a Mouse Model of Optogenetically Inducible Seizures. Neuron 2017, 93 (2), 291--298. 10.1016/j.neuron.2016.11.043.

(10) Molas, S.; Freels, T. G.; Zhao-Shea, R.; Lee, T.; Gimenez-Gomez, P.; Barbini, M.; Martin, G. E.; Tapper, A. R. Dopamine Control of Social Novelty Preference Is Constrained by an Interpeduncular-Tegmentum Circuit. Nat. Commun. 2024, 15 (1), 2891. 10.1038/s41467-024-47255-y.

(11) Formozov, A.; Dieter, A.; Wiegert, J. S. A Flexible and Versatile System for Multi-Color Fiber Photometry and Optogenetic Manipulation. *Cell Rep*. Methods 2023, 3 (3), 100418. 10.1016/j.crmeth.2023.100418.

(12) Sych, Y.; Chernysheva, M.; Sumanovski, L. T.; Helmchen, F. High-Density Multi-Fiber Photometry for Studying Large-Scale Brain Circuit Dynamics. Nat. Methods 2019, 16 (6), 553–560. 10.1038/s41592-019-0400-4.

(13) Osawa, S.; Iwasaki, M.; Hosaka, R.; Matsuzaka, Y.; Tomita, H.; Ishizuka, T.; Sugano, E.; Okumura, E.; Yawo, H.; Nakasato, N.; Tominaga, T.; Mushiake, H. Optogenetically Induced Seizure and the Longitudinal Hippocampal Network Dynamics. Plos One 2013, 8 (4), e60928. 10.1371/journal.pone.0060928.

(14) Klorig, D. C.; Alberto, G. E.; Smith, T.; Godwin, D. W. Optogenetically-Induced Population Discharge Threshold as a Sensitive Measure of Network Excitability. Eneuro 2019, 6 (6), ENEURO.0229--18.2019. 10.1523/eneuro.0229-18.2019.

(15) Gunaydin, L. A.; Grosenick, L.; Finkelstein, J. C.; Kauvar, I. V.; Fenno, L. E.; Adhikari, A.; Lammel, S.; Mirzabekov, J. J.; Airan, R. D.; Zalocusky, K. A.; Tye, K. M.; Anikeeva, P.; Malenka, R. C.; Deisseroth, K. Natural Neural Projection Dynamics Underlying Social Behavior. Cell 2014, 157 (7), 1535–1551. 10.1016/j.cell.2014.05.017.

(16) Akam, T.; Walton, M. E. PyPhotometry: Open Source Python Based Hardware and Software for Fiber Photometry Data Acquisition. Sci. Rep. 2019, 9 (1), 3521. 10.1038/s41598-019-39724-y.

(17) Lerner, T. N.; Shilyansky, C.; Davidson, T. J.; Evans, K. E.; Beier, K. T.; Zalocusky, K. A.; Crow, A. K.; Malenka, R. C.; Luo, L.; Tomer, R.; Deisseroth, K. Intact-Brain Analyses Reveal Distinct Information Carried by SNc Dopamine Subcircuits. Cell 2015, 162 (3), 635–647. 10.1016/j.cell.2015.07.014.

(18) Bruno, C. A.; O’Brien, C.; Bryant, S.; Mejaes, J.; Pizzano, C.; Estrin, D. J.; Barker, D. J. PMAT: An Open-Source, Modular Software Suite for the Analysis of Fiber Photometry Calcium Imaging. bioRxiv 2020, 2020.08.23.263673. 10.1101/2020.08.23.263673.

(19) Mueller, J.-S.; Tescarollo, F.; Huynh, T.; Brenner, D.; Valdivia, D.; Olagbegi, K.; Sangappa, S.; Chen, S.; Sun, H. Ictogenesis Proceeds Through Discrete Phases in Hippocampal CA1 Seizures. 2023. 10.21203/rs.3.rs-2613170/v1.

(20) Sarlo, G. L.; Holton, K. F. Brain Concentrations of Glutamate and GABA in Human Epilepsy: A Review. Seizure 2021, 91, 213–227. 10.1016/j.seizure.2021.06.028.

(21) Bruno, C. A.; O’Brien, C.; Bryant, S.; Mejaes, J. I.; Estrin, D. J.; Pizzano, C.; Barker, D. J. PMAT: An Open-Source Software Suite for the Analysis of Fiber Photometry Data. Pharmacol. Biochem. Behav. 2021, 201, 173093. 10.1016/j.pbb.2020.173093.

(22) Tescarollo; Valdivia; Chen, S.; Sun, H. Unilateral Optogenetic Kindling of Hippocampus Leads to More Severe Impairments of the Inhibitory Signaling in the Contralateral Hippocampus. Frontiers in Molecular Neuroscience 2024, No. 16. 10.3389/fnmol.2023.1268311.

(23) Vickstrom, C. R.; Snarrenberg, S. T.; Friedman, V.; Liu, Q. Application of Optogenetics and in Vivo Imaging Approaches for Elucidating the Neurobiology of Addiction. Mol. Psychiatry 2022, 27 (1), 640–651. 10.1038/s41380-021-01181-3.

(24) Lerner, T. N.; Shilyansky, C.; Davidson, T. J.; Evans, K. E.; Beier, K. T.; Zalocusky, K. A.; Crow, A. K.; Malenka, R. C.; Luo, L.; Tomer, R.; Deisseroth, K. Intact-Brain Analyses Reveal Distinct Information Carried by SNc Dopamine Subcircuits. Cell 2015, 162 (3), 635–647. 10.1016/j.cell.2015.07.014.

(25) Mejaes, J.; Desai, D.; Siciliano, C. A.; Barker, D. J. Practical Opinions for New Fiber Photometry Users to Obtain Rigorous Recordings and Avoid Pitfalls. Pharmacol. Biochem. Behav. 2022, 221, 173488. 10.1016/j.pbb.2022.173488.

(26) Qi, Z.; Guo, Q.; Wang, S.; Jia, M.; Gao, X.; Luo, M.; Fu, L.; China, B. C. C. for B. P., Wuhan National Laboratory for Optoelectronics, Huazhong University of Science and Technology, Wuhan 430074,; China, M. K. L. for B. P., School of Engineering Sciences, Huazhong University of Science and Technology, Wuhan 430074,; China, N. I. of B. S., Beijing 102206,; China, B. A. I. C. for B. D.-B. P. M., Beijing 100191,; China, S. of B. E., Capital Medical University, Beijing 100069,; China, C. I. for B. R., Beijing 102206,; China, S. of L. S., Tsinghua University, Beijing 100084,. All-Fiber-Transmission Photometry for Simultaneous Optogenetic Stimulation and Multi-Color Neuronal Activity Recording. Opto-Electron. Adv. 2022, 5 (12), 210081–210081. 10.29026/oea.2022.210081.

(27) Feng, J.; Zhang, C.; Lischinsky, J. E.; Jing, M.; Zhou, J.; Wang, H.; Zhang, Y.; Dong, A.; Wu, Z.; Wu, H.; Chen, W.; Zhang, P.; Zou, J.; Hires, S. A.; Zhu, J. J.; Cui, G.; Lin, D.; Du, J.; Li, Y. A Genetically Encoded Fluorescent Sensor for Rapid and Specific In Vivo Detection of Norepinephrine. Neuron 2019, 102 (4), 745–761.e8. 10.1016/j.neuron.2019.02.037.

(28) Chen, S. C.-Y.; Benvenuti, G.; Chen, Y.; Kumar, S.; Ramakrishnan, C.; Deisseroth, K.; Geisler, W. S.; Seidemann, E. Similar Neural and Perceptual Masking Effects of Low-Power Optogenetic Stimulation in Primate V1. eLife 2022, 11, e68393. 10.7554/elife.68393.

(29) Zong, W.; Obenhaus, H. A.; Skytøen, E. R.; Eneqvist, H.; Jong, N. L. de; Vale, R.; Jorge, M. R.; Moser, M.-B.; Moser, E. I. Large-Scale Two-Photon Calcium Imaging in Freely Moving Mice. Cell 2022, 185 (7), 1240–1256.e30. 10.1016/j.cell.2022.02.017.

(30) Sadakane, O.; Masamizu, Y.; Watakabe, A.; Terada, S.-I.; Ohtsuka, M.; Takaji, M.; Mizukami, H.; Ozawa, K.; Kawasaki, H.; Matsuzaki, M.; Yamamori, T. Long-Term Two-Photon Calcium Imaging of Neuronal Populations with Subcellular Resolution in Adult Non-Human Primates. Cell Rep. 2015, 13 (9), 1989–1999. 10.1016/j.celrep.2015.10.050.

(31) Cela, E.; Sjöström, P. J. A Step-by-Step Protocol for Optogenetic Kindling. Front Neural Circuit 2020, 14, 3. 10.3389/fncir.2020.00003.

(32) Shimoda, Y.; Beppu, K.; Ikoma, Y.; Morizawa, Y. M.; Zuguchi, S.; Hino, U.; Yano, R.; Sugiura, Y.; Moritoh, S.; Fukazawa, Y.; Suematsu, M.; Mushiake, H.; Nakasato, N.; Iwasaki, M.; Tanaka, K. F.; Tominaga, T.; Matsui, K. Optogenetic Stimulus-Triggered Acquisition of Seizure Resistance. Neurobiol Dis 2022, 163, 105602. 10.1016/j.nbd.2021.105602.

(33) Choy, M.; Dadgar-Kiani, E.; Cron, G. O.; Duffy, B. A.; Schmid, F.; Edelman, B. J.; Asaad, M.; Chan, R. W.; Vahdat, S.; Lee, J. H. Repeated Hippocampal Seizures Lead to Brain-Wide Reorganization of Circuits and Seizure Propagation Pathways. Neuron 2022, 110 (2), 221–236.e4. 10.1016/j.neuron.2021.10.010.

(34) Wang, Y.; Xu, C.; Xu, Z.; Ji, C.; Liang, J.; Wang, Y.; Chen, B.; Wu, X.; Gao, F.; Wang, S.; Guo, Y.; Li, X.; Luo, J.; Duan, S.; Chen, Z. Depolarized GABAergic Signaling in Subicular Microcircuits Mediates Generalized Seizure in Temporal Lobe Epilepsy. Neuron 2017, 95 (1), 92--105.e5. 10.1016/j.neuron.2017.06.004.

(35) Root, D. H.; Estrin, D. J.; Morales, M. Aversion or Salience Signaling by Ventral Tegmental Area Glutamate Neurons. iScience 2018, 2, 51–62. 10.1016/j.isci.2018.03.008.

(36) Millard, S. J.; Hoang, I. B.; Sherwood, S.; Taira, M.; Reyes, V.; Greer, Z.; O’Connor, S. L.; Wassum, K. M.; James, M. H.; Barker, D. J.; Sharpe, M. J. Cognitive Representations of Intracranial Self-Stimulation of Midbrain Dopamine Neurons Depend on Stimulation Frequency. Nat. Neurosci. 2024, 27 (7), 1253–1259. 10.1038/s41593-024-01643-1.

(37) Lévesque, M.; Avoli, M.; Kokaia, M. Jasper’s Basic Mechanisms of the Epilepsies. 2024, 236–240. 10.1093/med/9780197549469.003.0011.

(38) Bansal, H.; Gupta, N.; Roy, S. Comparison of Low-Power, High-Frequency and Temporally Precise Optogenetic Inhibition of Spiking in NpHR, ENpHR3.0 and Jaws-Expressing Neurons. Biomed. Phys. Eng. Express 2020, 6 (4), 045011. 10.1088/2057-1976/ab90a1.

(39) Lin, J. Y. A User’s Guide to Channelrhodopsin Variants: Features, Limitations and Future Developments. Exp. Physiol. 2010, 96 (1), 19–25. 10.1113/expphysiol.2009.051961.

(40) Nakagawa, T.; Oghalai, J. S.; Saggau, P.; Rabbitt, R. D.; Brownell, W. E. Photometric Recording of Transmembrane Potential in Outer Hair Cells. J. Neural Eng. 2006, 3 (2), 79. 10.1088/1741-2560/3/2/001.

(41) Sahu, A.; Bhat, V. N.; Patra, S.; Tiwari, V. High-Sensitivity Fluorescence-Detected Multidimensional Electronic Spectroscopy through Continuous Pump–Probe Delay Scan. J. Chem. Phys. 2023, 158 (2), 024201. 10.1063/5.0130887.

(42) Dorsa, A.; Xie, Q.; Wagner, M.; Xu, X. G. Lock-in Amplifier Based Peak Force Infrared Microscopy. Anal. 2023, 148 (2), 227–232. 10.1039/d2an01103d.

(43) Seto, K.; Okuda, Y.; Tokunaga, E.; Kobayashi, T. Development of a Multiplex Stimulated Raman Microscope for Spectral Imaging through Multi-Channel Lock-in Detection. Rev. Sci. Instrum. 2013, 84 (8), 083705. 10.1063/1.4818670.

(44) Yan, Y.; Petchprayoon, C.; Mao, S.; Marriott, G. Reversible Optical Control of Cyanine Fluorescence in Fixed and Living Cells: Optical Lock-in Detection Immunofluorescence Imaging Microscopy. Philos. Trans. R. Soc. B: Biol. Sci. 2013, 368 (1611), 20120031. 10.1098/rstb.2012.0031.

(45) Zhang, X.; Zhai, D.; Li, T.; Zhou, Y.; Lin, Y. Image Inpainting Based on Deep Learning: A Review. Inf. Fusion 2023, 90, 74–94. 10.1016/j.inffus.2022.08.033.

(46) Li, P.; Geng, X.; Jiang, H.; Caccavano, A.; Vicini, S.; Wu, J. Measuring Sharp Waves and Oscillatory Population Activity With the Genetically Encoded Calcium Indicator GCaMP6f. Front. Cell. Neurosci. 2019, 13, 274. 10.3389/fncel.2019.00274.

(47) Kim, C. K.; Yang, S. J.; Pichamoorthy, N.; Young, N. P.; Kauvar, I.; Jennings, J. H.; Lerner, T. N.; Berndt, A.; Lee, S. Y.; Ramakrishnan, C.; Davidson, T. J.; Inoue, M.; Bito, H.; Deisseroth, K. Simultaneous Fast Measurement of Circuit Dynamics at Multiple Sites across the Mammalian Brain. Nat. Methods 2016, 13 (4), 325–328. 10.1038/nmeth.3770.

(48) Soor, N. S.; Quicke, P.; Howe, C. L.; Pang, K. T.; Neil, M. A. A.; Schultz, S. R.; Foust, A. J. All-Optical Crosstalk-Free Manipulation and Readout of Chronos-Expressing Neurons. J. Phys. D 2019, 52 (10), 104002. 10.1088/1361-6463/aaf944.

(49) Guo, Q.; Zhou, J.; Feng, Q.; Lin, R.; Gong, H.; Luo, Q.; Zeng, S.; Luo, M.; Fu, L. Multi- Channel Fiber Photometry for Population Neuronal Activity Recording. Biomed. Opt. Express 2015, 6 (10), 3919–3931. 10.1364/boe.6.003919.

