## Supplemental Figures for "µFIX – Enabling Combinations of Concurrent Optogenetics and Lock-in Amplification Fiber Photometry via Removal of Optogenetic Stimulation Crosstalk"

Supporting Information

| **Content** | Page |
| --- | --- |
| Figure S1. Additional examples of before-and-after crosstalk removal. | S2 |
| Figure S2. Power spectral density charts of empirical and simulated recordings. | S3 |
| Figure S3. Recovery fidelity vs. variability of source signal. | S4 |
| Figure S4. Recovery results using smaller amounts of remaining data for fitting. | S5 |
| Figure S5. Recovery results at 30, 60, and 90 s stimulation length. | S6 |

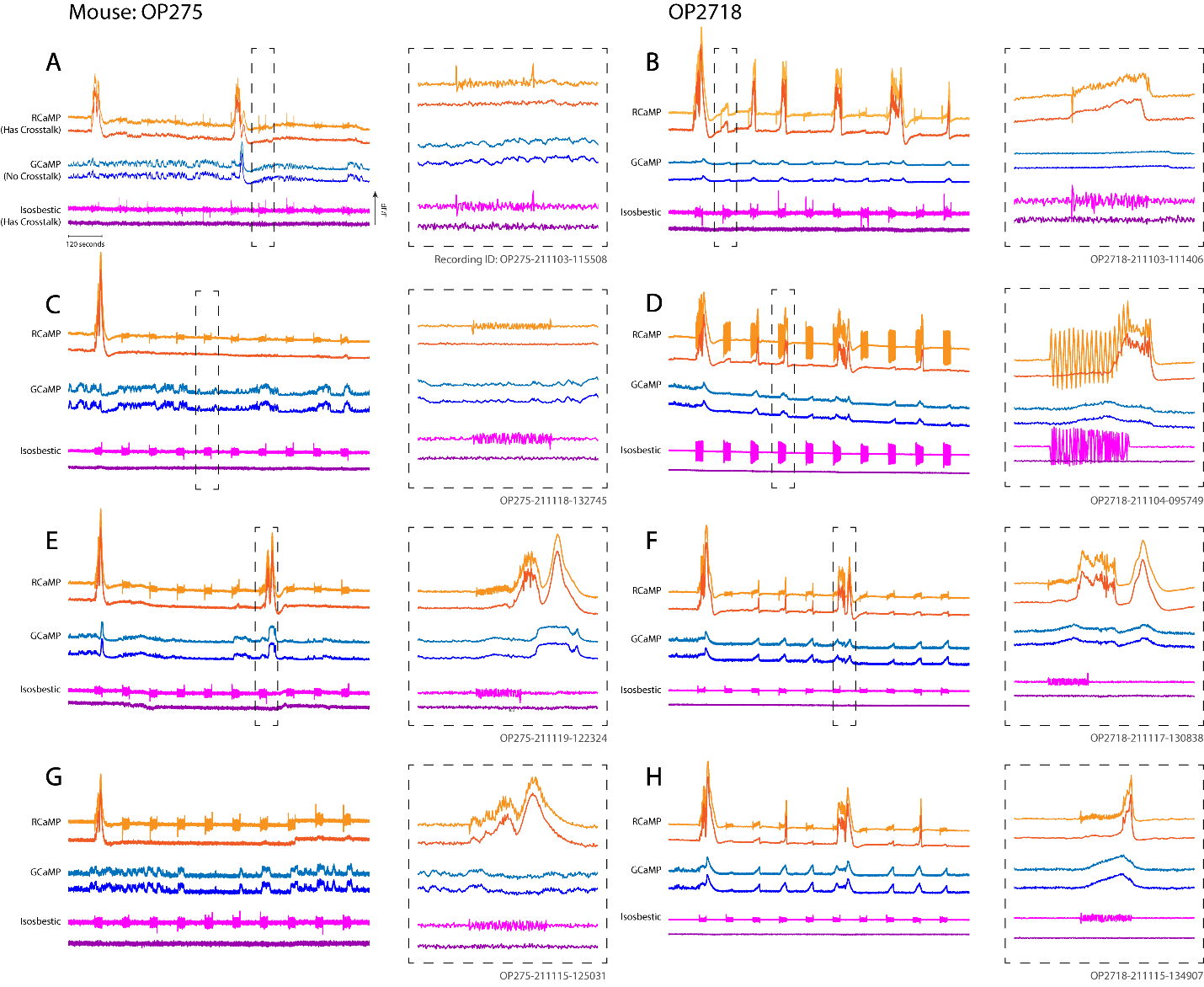

**Figure S1.** *Additional examples of recordings before and after crosstalk removal via µFIX from Mouse OP275 at left and Mouse OP2718 at right. Dashed boxes show zoomed-in single epochs for detail. A, B, C, and H zoomed-in sections show non-seizure responses and D, E, F, and G show a seizure response.*

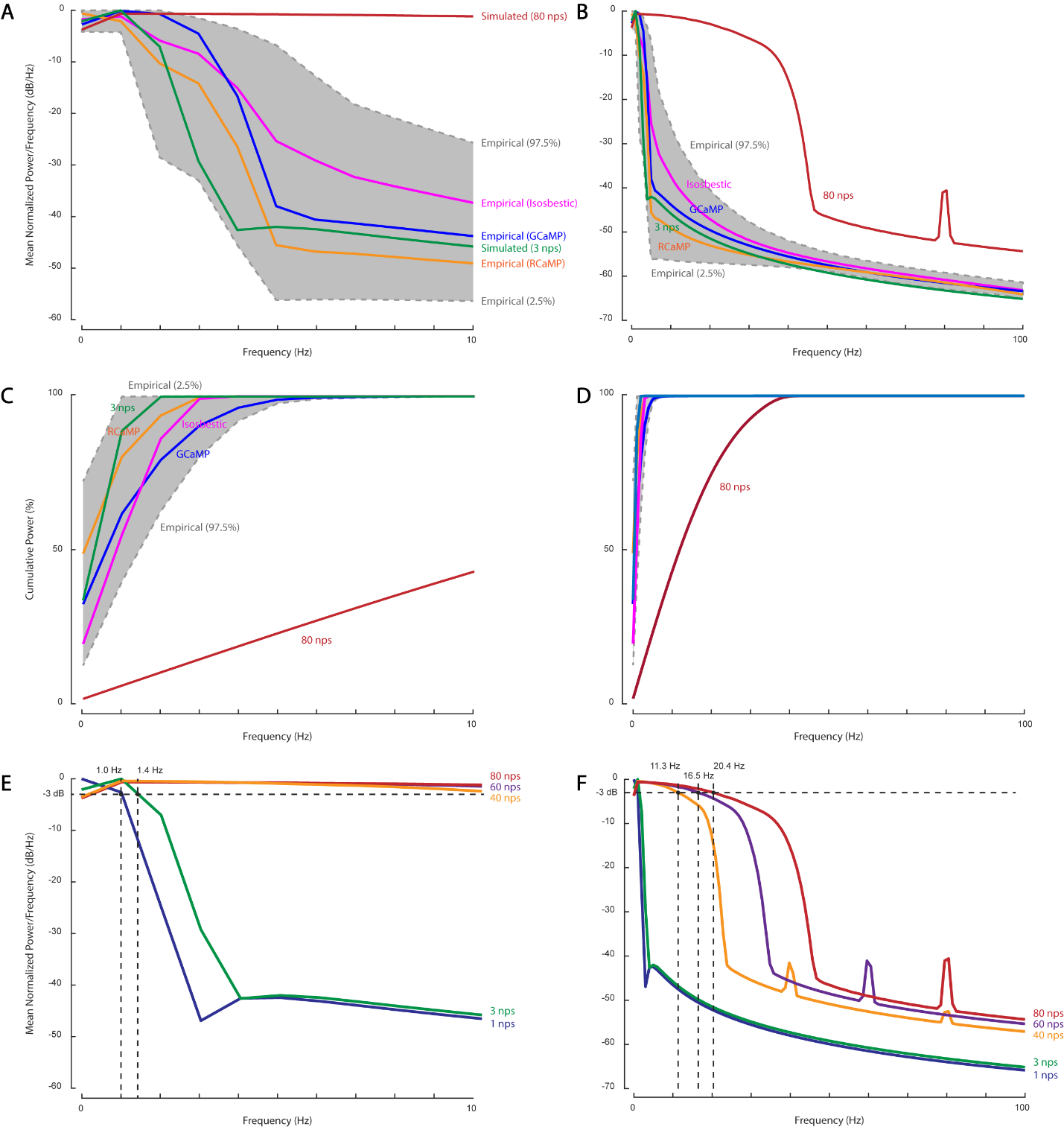

**Figure S2.** *Comparison of power spectral densities between empirical recordings, 3 and 80 nps simulated recordings.* ***(A-B)*** *Mean normalized power (n = 455 in empirical epochs, n = 600 in simulated epochs), computed by the pwelch MATLAB function on each epoch, then converted to logarithmic decibel scale, normalized to the maximum power in each epoch, and then averaged across epochs. The shaded area shows the 95% confidence interval of the combined empirical data power spectrum.* ***(C-D)*** *Normalized cumulative power of (A-B).* ***(E-F)*** *Mean normalized power as in (A-B) for additional nps values. The limit of the frequency content represented with each nps value is estimated at -3 dB.*

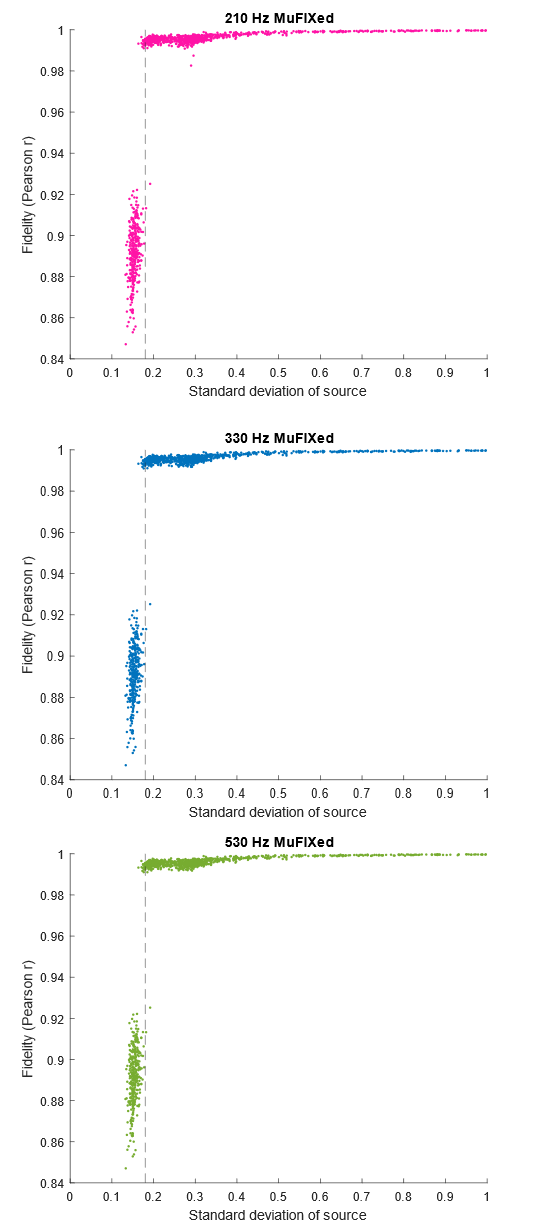

**Figure S3.** *Plot of recovery fidelity for empirical ground truth signals encoded on all three carrier frequencies (CT+µFIX) against the standard deviation of the source signal. µFIX had a clear minimum SD at around 0.18 mV (marked by the vertical dashed line) to produce optimal signal recovery from crosstalk.*

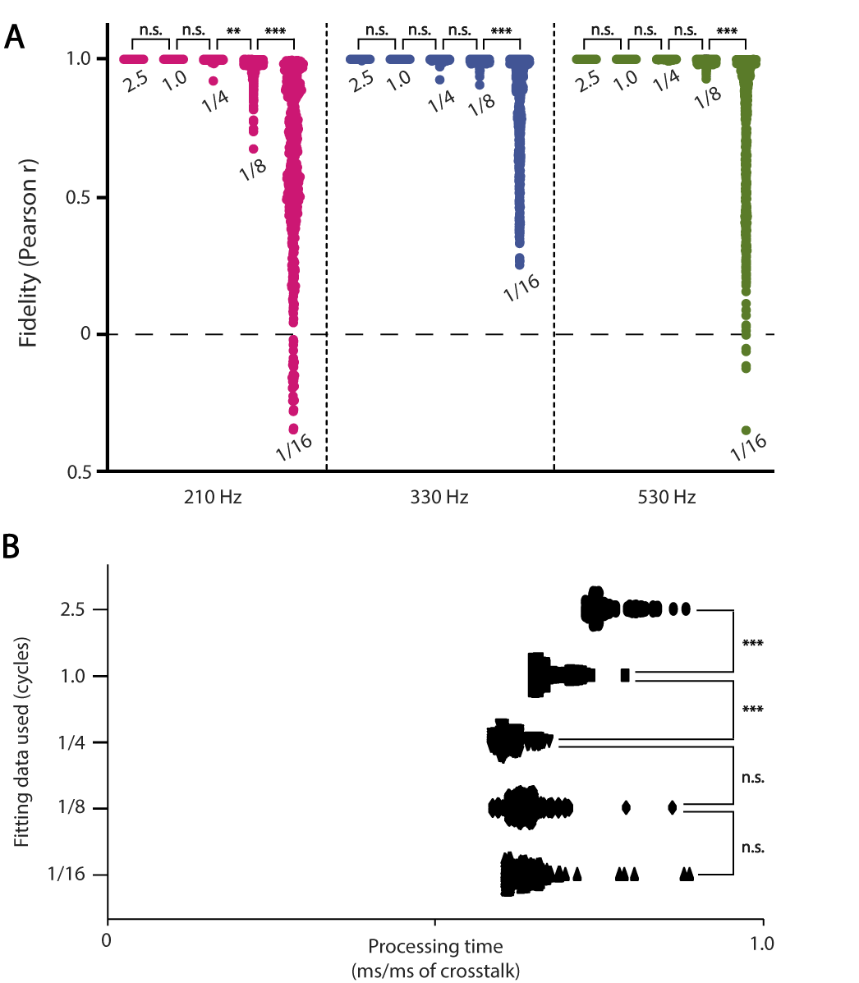

**Figure S4. *(A)*** *Fidelity of crosstalk removal with respect to the length of intact data before and after crosstalk. The length of intact data is labeled for each column, expressed in cycles (1/16 to 2.5 cycles) of 210 Hz, the lowest encoding frequency. Simulations were run with artificial ground truth signals generated at 3 nps, and simulated optogenetic stimulation of 20 Hz 5 ms pulses for 30 s.* ***(B)*** *Processing time with respect to the length of intact data used. Results are shown in milliseconds of processing time per millisecond of crosstalk (i.e. 5 ms stim + 4ms artefact = 9 ms of crosstalk per pulse). N = 600 trials for every cycle length. T-tests: n.s. = not significant, * p < 0.05, ** p < 0.01, *** p < 0.001.*

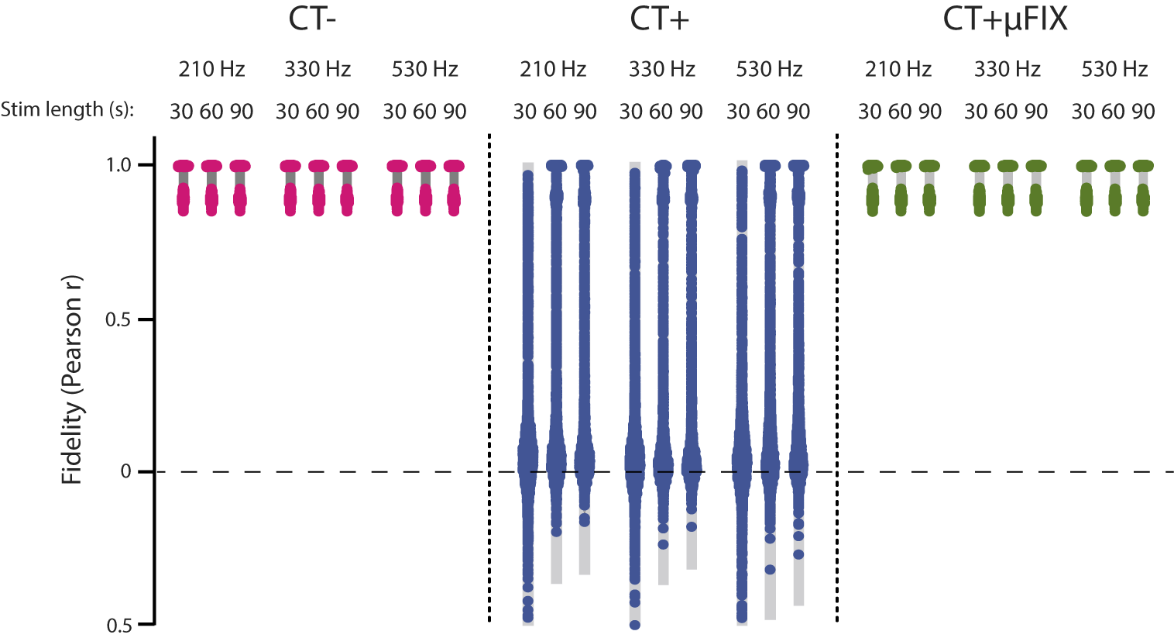

**Figure S5.** *Fidelity of crosstalk signal recovery with respect to the length of stimulation train. The length of the stimulation train (30, 60, and 90 s) is labeled for each column. The 30 s results are the same as reported in Fig. 6. For 60 and 90 s, the same empirical ground truth recordings were used with the same stimulation protocol of 5 ms pulses at 20 Hz. The fidelity (Pearson correlation) calculation window was expanded for 60 and 90 s, respectively.*
